# Genomic islands of speciation harbor genes underlying coloration differences in a pair of Neotropical seedeaters

**DOI:** 10.1101/2023.11.14.567069

**Authors:** Tram N. Nguyen, Márcio Repenning, Carla Suertegaray Fontana, Leonardo Campagna

**Affiliations:** Department of Ecology and Evolutionary Biology, Cornell University, 215 Tower Road, Ithaca, NY 14853, USA; Universidade Federal do Rio Grande (FURG), Laboratório de Aves Aquáticas e Tartarugas Marinhas (LAATM). Av. Itália km 8, Campus Carreiros, 96203-900, Rio Grande, Rio Grande do Sul, Brazil; Universidade Federal do Rio Grande do Sul (UFRGS). Laboratório de Ecologia de Comunidades e Populações, Instituto de Biociências, Avenida Bento Gonçalves, 9500, 91501-970, Porto Alegre, Rio Grande do Sul, Brazil; Programa de Pós-graduação em Biodiversidade Animal, Universidade Federal de Santa Maria.; Fuller Evolutionary Biology Program, Cornell Lab of Ornithology, 159 Sapsucker Woods Road, Ithaca, NY, 14850, USA

**Keywords:** Ancestral Recombination Graph, Genome scan, Hybridization, Islands of speciation, *Sporophila*

## Abstract

Incomplete speciation can be leveraged to associate phenotypes with genotypes, thus providing insights into the traits relevant to the reproductive isolation of diverging taxa. We investigate the genetic underpinnings of the phenotypic differences between *Sporophila plumbea* and *S. beltoni*. *S. beltoni* has only recently been described based, most notably, on differences in bill coloration (yellow vs. black in *S. plumbea*). Both species are indistinguishable through mtDNA or reduced-representation genomic data, and even whole-genome sequencing revealed low genetic differentiation. Demographic reconstructions attribute this genetic homogeneity to gene flow, despite divergence in the order of millions of generations. We found a narrow hybrid zone in southern Brazil where genetically, yet not phenotypically, admixed individuals appear to be prevalent. Despite the overall low genetic differentiation, we identified three narrow peaks along the genome with highly differentiated SNPs. These regions harbor six genes, one of which is involved in pigmentation (*EDN3*) and is a candidate for controlling bill color. Within the outlier peaks we found signatures of resistance to gene flow, as expected for islands of speciation. Our study shows how genes related to coloration traits are likely involved in generating prezygotic isolation and establishing species boundaries early in speciation.

## Introduction

Increasing knowledge about the genetic, ecological, and biogeographic contexts in which new species are formed is central to understanding the processes which lead to speciation. General patterns can be found by accumulating examples of how speciation has taken place throughout the tree of life [1–4]. The genetic architecture of reproductive barriers can be studied by leveraging non-model systems in the early stages of speciation and/or by focusing on hybridizing species where these barriers break down. For recently diverged taxa, genetic differentiation tends to be localized around areas relevant to speciation, but with time, genomes will continue to diverge as other differences start to follow [5]. In such cases, natural hybridization or experimental crossing can help break up genetic linkage through recombination, and phenotypic traits can still be associated statistically to genetic changes [6]. Whereas studies following these types of designs have revealed how genomic landscapes are clearly heterogeneously differentiated [5], the processes driving such differentiation remain debated. Areas of high differentiation (or outlier regions) can be shaped by processes such as demographic range expansions [7], background selection combined with low levels of recombination [8], selective sweeps [9] or resistance to gene flow [10,11]. These processes can leave different signatures in the outlier genomic regions which researchers can examine with various traditional summary statistics, such as F_ST_, D_XY_, Tajima’s D, and Pi, to uncover why genomes are heterogeneously differentiated [11]. However, clarity on the processes which have shaped many of these genomic regions is still generally lacking [8]. With the increasing feasibility of whole-genome resequencing, the toolkit for inferring evolutionary processes from genomic patterns continues to grow, offering researchers greater power to pinpoint regions of the genome under selection that mediate phenotypes of interest [12].

In birds, genome scans among recently diverged taxa have repeatedly uncovered coloration genes in areas of high differentiation in species that otherwise show shallow overall levels of genomic differentiation [10,13–17]. Collectively, these findings point to the importance of coloration differences in the early stages of speciation, and their likely effectiveness as mechanisms of prezygotic isolation [18–20]. Notably, *de novo* mutations in coloration genes, or the reassembly of different variants from standing variation across permeable species limits [21], may promote speciation. However, many taxa in the early stages of speciation may be ephemeral [22,23], with their persistence dependent on the accumulation of additional mutations over time, perhaps facilitated by the biogeographic context in which they were formed. Nevertheless, recently diverged species, whether they persist or not, will be informative of the types and location of genetic changes which can lead to initiating speciation.

The Neotropical genus *Sporophila* contains over 40 species and has the highest speciation rates of the species-rich Tanager family, to which it belongs [24,25]. This high speciation rate is in part the consequence of several small groups of rapidly radiating (recent and closely related) taxa in the early stages of speciation, making the genus a valuable study system to understand the origin of reproductive isolation and the process of speciation. Many of these groups have been studied from a genetic perspective, and consistently show differentiation in male sexual traits (including coloration patterns) despite little genome-wide genetic differentiation [14,26,27]. The largest of such groups is known as the capuchino seedeaters, which contains 12 species differing most notably in the plumage patterns of adult males and in their vocalizations (a primarily cultural trait) [28,29]. Despite their phenotypic diversity, capuchino seedeaters show extremely low genetic differentiation, except for a few areas of the genome which are enriched for pigmentation genes. These genes likely mediate the different coloration phenotypes observed across males from different species [14,20,30], which in turn, impact species recognition [20]. Similarly, the Variable Seedeater superspecies complex (*S. corvina*, *S. intermedia*, *S. murallae*, and *S. americana*) and Morelet’s Seedeater (*S. morelletti*) represent additional cases of plumage diversity, yet shallow genetic differentiation within *Sporophila* [26,27,31,32]. Overall, these examples suggest that localized genetic differences may mediate divergence in coloration traits, contributing to reproductive isolation and speciation [20].

Another *Sporophila* species that is not part of the previously mentioned groups, the Plumbeous Seedeater (*S. plumbea*), was traditionally thought to contain black and yellow-billed individuals. However, this presumed intraspecific variation was not a common polymorphism and did not correlate with seasonality or age (Figure 1A and 1B, [33]). In 2013, the yellow-billed individuals were shown to constitute a new species, the Tropeiro seedeater (*S. beltoni*), that bred in the grassland highlands of southern Brazil [34,35]. *S. beltoni* is differentiated from *S. plumbea* primarily through bill coloration, but also through male size and bill shape, vocalizations, migration phenology and subtle aspects of male plumage coloration. Furthermore, both species also differ in their breeding ranges, habitat, and phenology [34,35]. *S. beltoni* is sexually dimorphic and shares a narrow contact zone (∼50-100 km) in the north of its distribution with the southernmost breeding population of *S. plumbea* (Figure 1A). In this contact zone, individuals are mostly segregated by habitat and elevation, though they can breed in close proximity where their respective habitats are available [34]. Although hybrids have not been definitively identified in this region, a few individuals with irregularly colored and streaky bills were observed, raising the possibility that hybridization occurs in this contact zone. However, a preliminary study using reduced-representation genomic data failed to distinguish these two species [35], suggesting they were of very recent origin and/or continued to experience high levels of gene flow. Together, the phenotypic difference and low genetic differentiation between this species pair provides an opportunity to further investigate the process of speciation.

**Figure 1:**
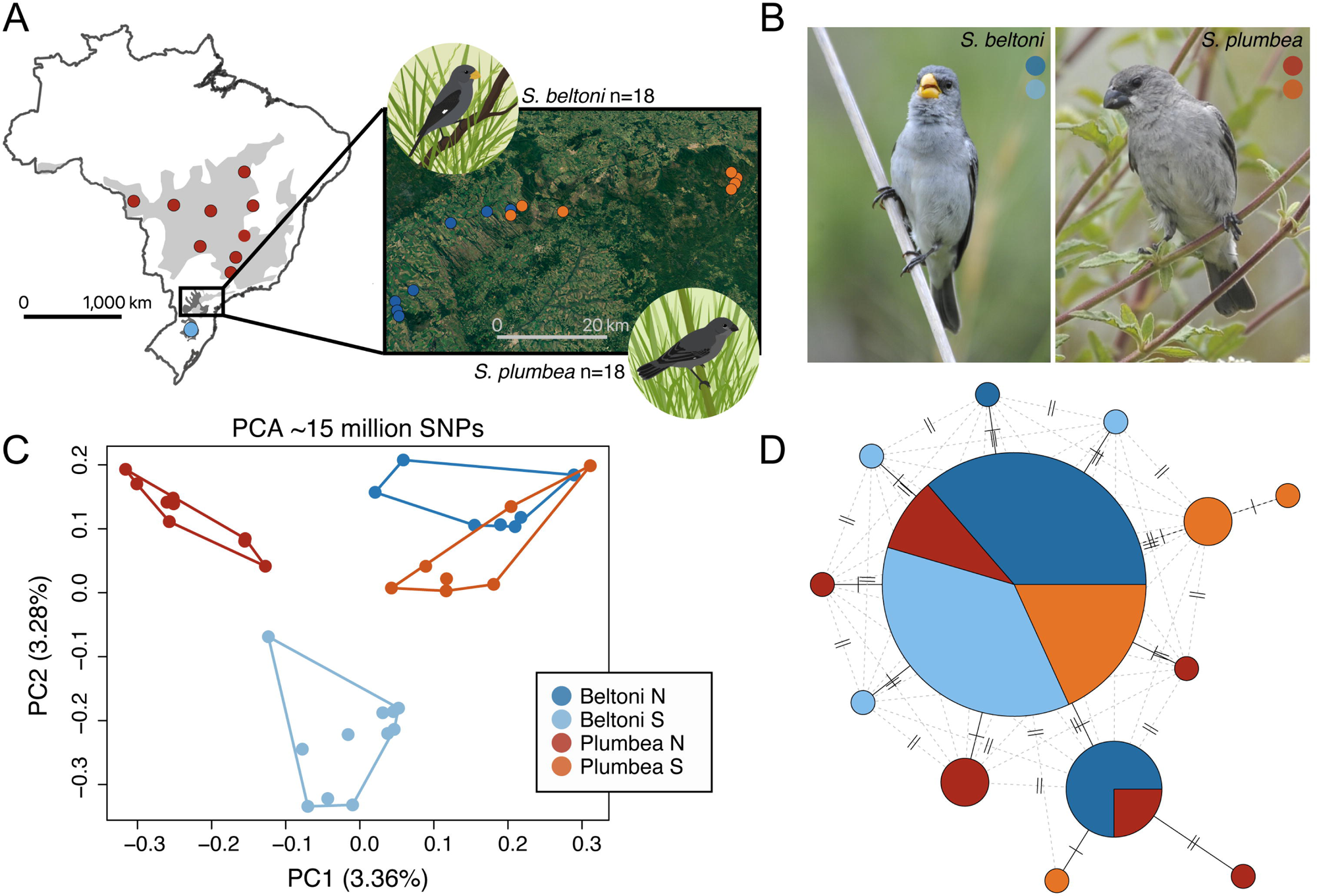
Genome-wide differentiation between *S. beltoni* and *S. plumbea*. **(A)** Range map for both species showing sampling localities in Brazil, and the area of contact (encompassed in the rectangle on the map and also in the topographic map in the inset) following [34]. Samples are color-coded by species and geographic origin (as shown in **C**). (**B**) Examples of adult male *S. beltoni* and *S. plumbea* individuals; photos by Márcio Repenning. (**C**) PCA derived from ∼15 million genome-wide SNPs. (**D**) Haplotype network derived from sequences of the mitochondrial gene *COI*. Different haplotypes are connected either by solid (based on the minimum spanning tree) or by dashed lines (indicating alternative paths). The number of mutational steps between haplotypes is shown as solid bars.

Here we assessed the degree of genetic differentiation between *S. beltoni* and *S. plumbea* using whole-genome resequencing, and conducted the first study to search for genomic regions associated with their phenotypic differences. Using the genomes of individuals from each species across both their allopatric ranges (i.e., outside of the contact zone) and within the contact zone, we conducted a demographic reconstruction to understand the demographic patterns in these different geographic regions. We then performed a genome scan which highlighted three narrow regions containing genes that likely mediate differences in beak coloration. We subsequently used genealogy-based statistics derived from topologies extracted from the ancestral recombination graph (ARG) to infer the processes that have shaped genomic outlier regions, and found signatures of reduced gene flow, as is expected for islands of speciation. Our study contributes to the emerging body of literature showing differentiation in genes related to bird coloration, which may mediate prezygotic isolation, before speciation is complete.

## Results

### Low genomic differentiation between *S. beltoni* and *S. plumbea*

While *S. beltoni* and *S. plumbea* individuals sampled outside of the contact zone (n=11 for each species) clustered separately in a PCA derived from ∼15 million genome-wide SNPs, those sampled within the contact zone resembled each other, overlapping in the PCA (n=7 for each species; Figure 1C, Figure S1). Moreover, individuals could not be assigned to species using mtDNA (cytochrome c oxidase subunit 1 sequences), irrespective of sampling location (Figure 1D). Although the overall level of genome-wide differentiation was consistently low across groups, it was higher when comparing between allopatric individuals versus between sympatric individuals (F_ST_ = 0.0031 for allopatric, ∼0 for sympatric, and on average 0.0015 for all samples combined). Using these genomic data, we then reconstructed the history of divergence between the two species. We found similarly large current effective population sizes for both taxa (in the order of millions of individuals), which were much larger than the size of the inferred ancestral population (∼5 fold; Figure 2A). The divergence time between *S. beltoni* and *S. plumbea* was estimated to be between ∼3.2 and 3.6 million generations (Figure 2B), and the genetic similarity between the species could be partly explained by high levels of gene flow since their split (between 2 and 4 migrants per generation; Figure 2C). There was some variation in the parameter values obtained from models using birds sampled from different parts of the range (allopatric, sympatric, or a combination of both; Figure 2). However, the most notable difference was the direction of gene flow (Figure 2C), which we interpret with caution as it can be particularly challenging to infer under certain scenarios [36]. While migration was inferred from *S. beltoni* into *S. plumbea* within the contact zone, the opposite was true for allopatric individuals (or when all samples were combined). It is possible that the patterns of hybridization may be different within and outside of the contact zone, with the former being representative of more recent gene flow (see Discussion).

**Figure 2:**
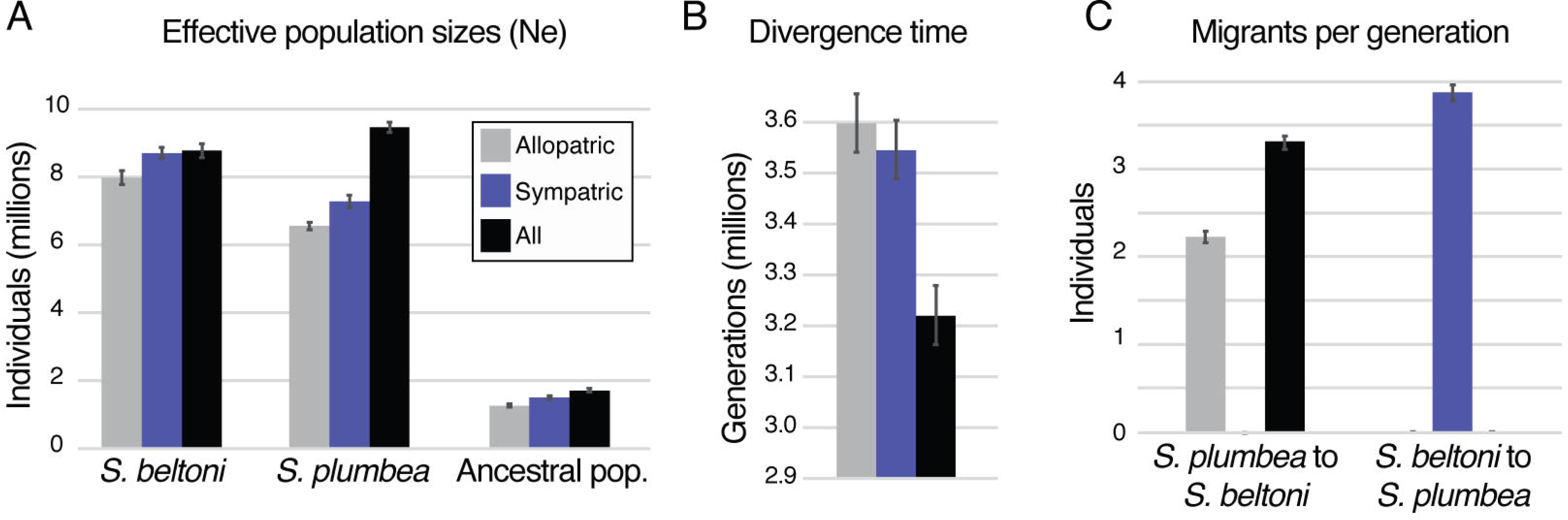
Demographic reconstruction. (**A**) Current and ancestral population sizes, (**B**) divergence time and (**C**) migrants per generation. Each set of parameters was estimated three times, once with a random subset of all samples combined and also after subsampling either exclusively allopatric or sympatric individuals. Bars indicate median values and 95% credible intervals.

### Differentiated SNPs are concentrated in three outlier peaks

When comparing all samples, a small number of SNPs showed differentiation above F_ST_=0.20 (1.1%) and only 40 SNPs (out of ∼15 million) had an F_ST_ value above 0.7 (Figure 3A). Of the 40 variants showing the highest F_ST_ values, 39 were clustered in three divergence peaks: on contigs 404 (24 elevated SNPs in ∼100kb on the Z chromosome), 33 (9 elevated SNPs in ∼66 kb on chromosome 20), and 382 (6 elevated SNPs in ∼2 kb on chromosome 11) (Figure 3A). Taken together, the variants from these peaks could be combined to cluster individuals by species in a PCA (Figure S1), and the same was true when distinguishing allopatric individuals using the variants from each peak separately (Figure 3B). However, with the exception of contig 33, sympatric individuals were genetically similar to each other and could not always be assigned to species based on their genotypes in these genomic regions (Figure 3B, Figure S2). The 39 variants showing the highest F_ST_ values in these peaks were in high linkage-disequilibrium, both within but also between peaks, suggesting they are co-inherited despite being on different chromosomes (Figure S3). Moreover, each species showed a common haplotype on each peak, with less intraspecific variation compared to interspecific differences (Figures S4-S6). Only a few sympatric individuals were heterozygotes or sometimes homozygotes for haplotypes from the other species. We did not find individuals that showed admixture at all three peaks simultaneously (Figures S4-S6).

**Figure 3:**
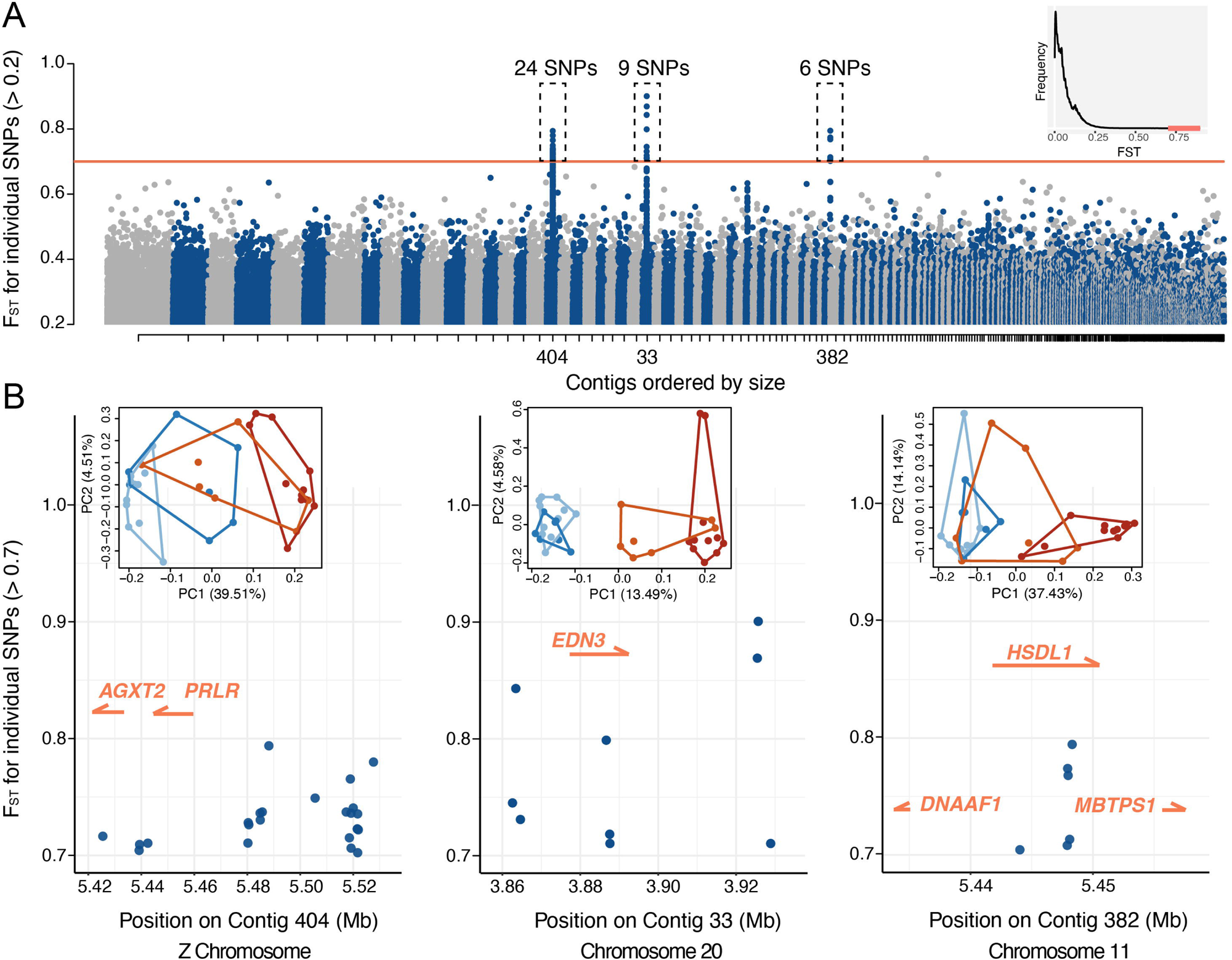
Outlier peaks between *S. beltoni* and *S. plumbea*. (**A**) Manhattan plot showing F_ST_ values calculated using all individuals from both species. Note that for simplicity the plot displays the 1.1% of SNPs with F_ST_ > 0.2. The inset shows a histogram of the distribution of F_ST_ values from all genome-wide SNPs, with those showing values above 0.7 in red. The plots in (**B**) represent a zoom into each peak shown in (**A**) and display the genes that are annotated in that region. The insets show PCAs derived from the variants found in each peak. Samples are color-coded by species and the geographic region to which they belong (as in Figure 1).

### *EDN3* and *PRLR* are candidate genes implicated in the phenotypic differences between species

The divergence peaks we identified involved a total of six genes (Figure 3B, Table 1). Two of these genes, *EDN3* which encodes the protein Endothelin-3, and *PRLR* which encodes the Prolactin receptor, may mediate the difference in beak coloration between *S. beltoni* and *S. plumbea* (Table 1; see Discussion). The remaining genes could be related to differences between these taxa that are not immediately obvious to us (Table 1). The variants showing the highest F_ST_ values in these peak regions were mostly non-coding (74% fell in intergenic regions and 13% in introns; Figure 3B), however two were responsible for non-synonymous mutations in the *HSDL1* gene on contig 382. Three additional mutations fell within *EDN3* but our annotation of that region was not of sufficient quality to distinguish if they fell within introns or in exons.

**Table 1.**
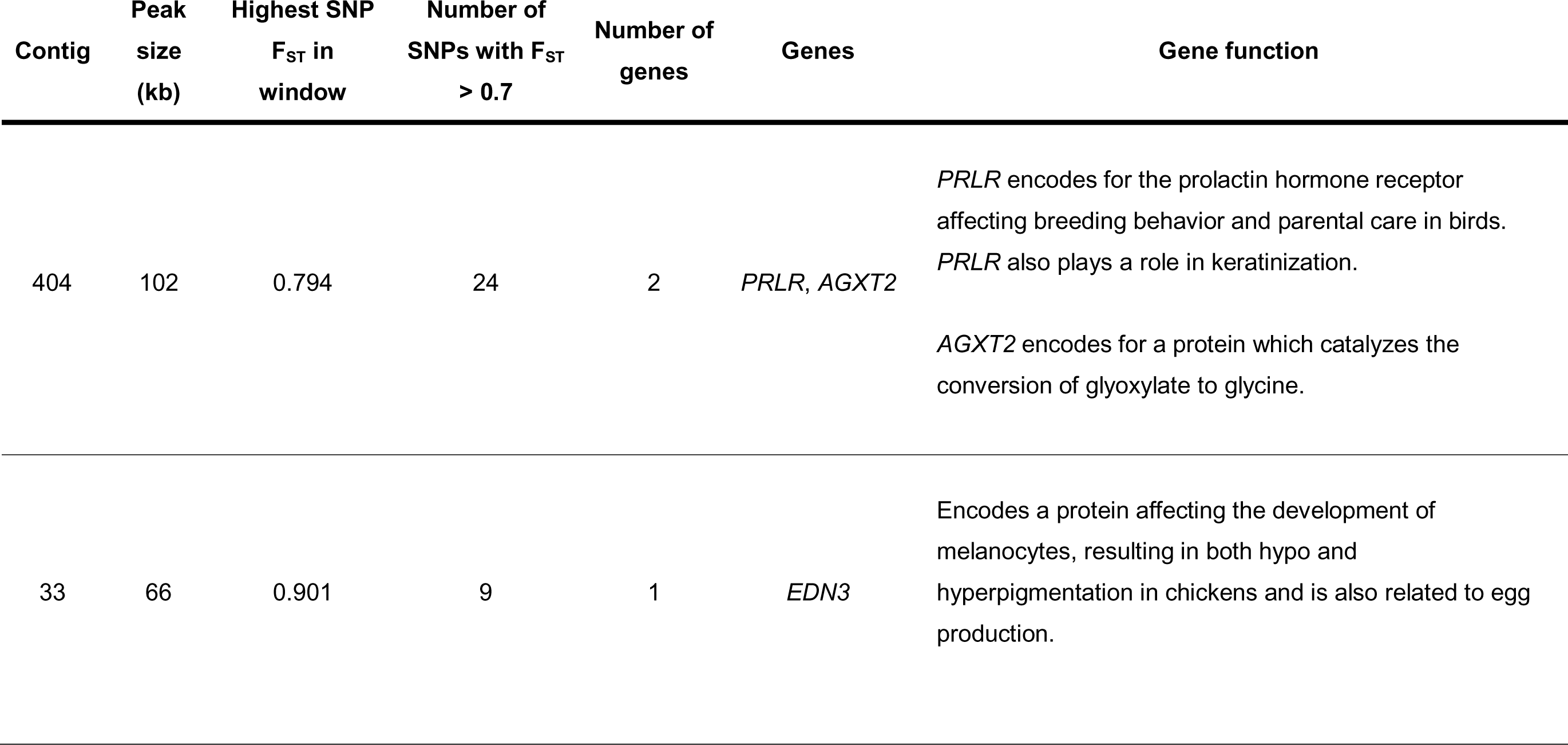
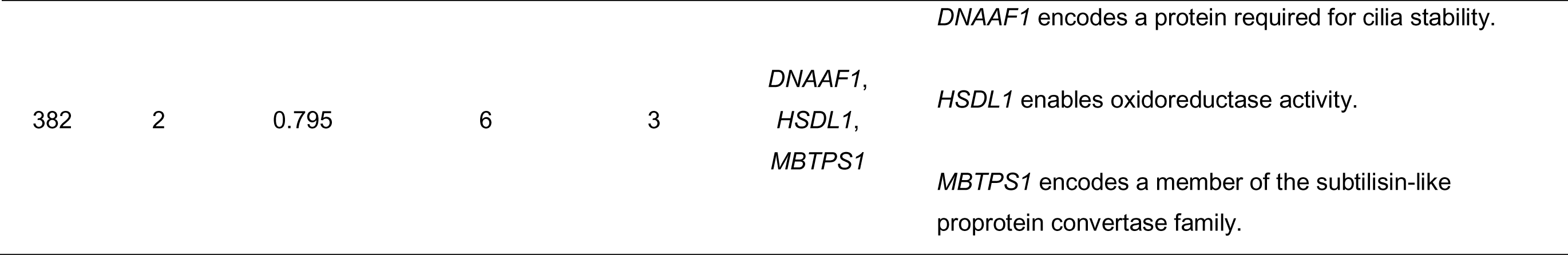
Regions of high divergences between species. Details for each outlier peak, including the genes they contain and a summary of their functions.

### Outlier genomic regions are speciation islands

We explored the divergence peaks to search for signatures that could reveal the processes involved in shaping these regions. We first looked at Tajima’s D and nucleotide diversity, which tend to be reduced after selective sweeps. Although Tajima’s D was on average negative and nucleotide diversity was reduced in the three peaks, more extreme values could be found outside of the peaks within the same contigs (Figure S7). Therefore, based on these statistics alone, it is not conclusive whether selective sweeps have occurred within these outlier peaks. We next looked at H12 and H2/H1 which can be used to distinguish hard and soft selective sweeps, but again did not find conclusive patterns (Figure S7). We also compared F_ST_ and D_XY_ values and observed slightly elevated D_XY_ in some windows within the F_ST_ divergence peaks (compared to both windows outside of the F_ST_ peaks or in contigs without F_ST_ peaks), suggesting these regions may have accumulated some absolute sequence differences between the species (Figure S8). This prompted us to search for signatures of selection in more depth using three statistics derived from topologies extracted from the ancestral recombination graph (ARG) (Figure 4A). The species enrichment score measures the probability of observing clades containing individuals from a certain species [9]. The relative TMRCA half-life, RTH’ [37], is a normalized version of the time to most recent common ancestry (TMRCA). Finally, the inter-species cross-coalescence measures the average age of the most recent coalescence events between tips in the tree which belong to the different species. We followed the reasoning outlined by Hejase et al. [9], where areas under selection are expected to show species enrichment, but these could be the product of either species-specific and recent selective sweeps, or they could be older islands of speciation that resist gene flow. These two extreme scenarios can be distinguished by using RTH’, which we expect to be low when selective sweeps produce shallow clades, and the inter-species cross-coalescence, which we expect to show larger values when gene flow is selected against (i.e., delayed cross-coalescence [9]). Following this logic, we obtained values for the three statistics in the divergence peaks and for a control set of contigs which represented ∼10% of the genome, and we used the latter to establish species-specific thresholds of significance for each statistic. We also conducted this analysis for all samples together, and for sympatric and allopatric samples separately. The analysis shows genealogies within the peak regions which are significantly enriched for each species (Figure 4B, Table 2, Table S2, Figures S9-S12). These clades show low RTH’ values, sometimes for one species but in some cases for both, and for the peaks on contigs 404 and 382 we also observed delayed cross-coalescence (Table 2, Table S2). These patterns were accentuated when conducting the analysis with samples from outside of the contact zone and generally weaker or lost when using sympatric birds from within the contact zone. Taken together, these regions show evidence of having undergone selection and experiencing reduced migration compared to other areas in the genome, especially when analyzing allopatric samples which may be more representative of historical processes (see Discussion).

**Figure 4:**
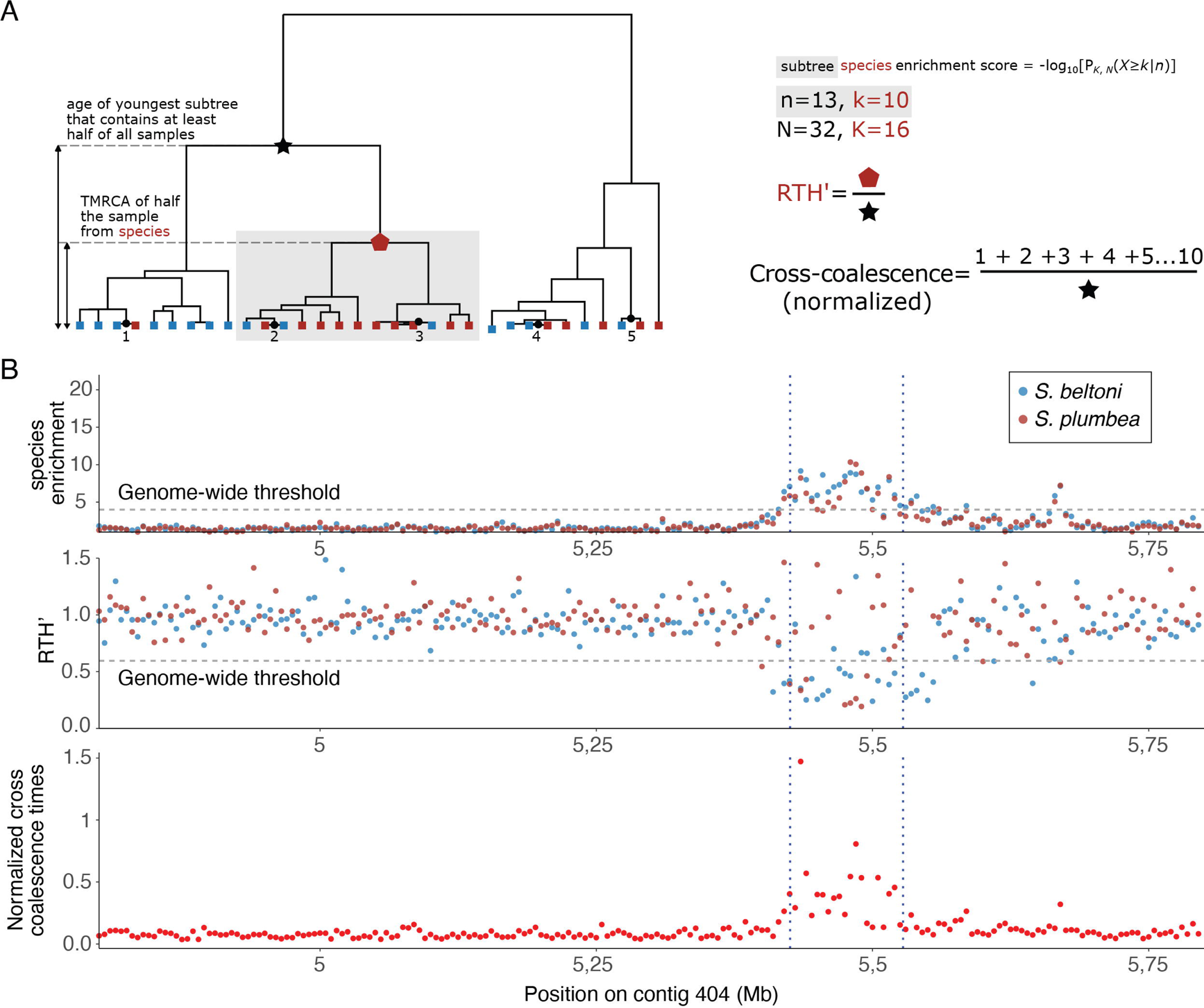
ARG-based statistics. (**A**) Graphical representation of how species enrichments, RTH’ and cross-coalescence values are calculated (modified from [17]). The species enrichment score is calculated for each subtree (example shaded in gray) in a topology and represents the probability of observing that number of samples of a particular species (the one represented in red in the example) in that subtree under a hypergeometric distribution. The score for a given tree and species is the maximum score associated with the full tree. RTH’ is the ratio between the TMRCA of half of the samples from a species and the age of the youngest subtree that contains at least half of all samples (irrespective of species). The cross-coalescence time is calculated by adding the time of the ten most recent cross-coalescence events in the tree and dividing it by the same value used to normalize TMRCA to obtain RTH’. In the example the color of the leaves indicates two hypothetical species. (**B**) Plots showing these statistics for contig 404 in 5 kb windows, with the peak region shown by dashed vertical blue lines. These particular plots show statistics derived from individuals sampled outside of the hybrid zone. Species enrichment values for *S. beltoni* and *S. plumbea* are higher than the genome-wide threshold of statistical significance, while RTH’ values are significantly lower. Therefore, subtrees from topologies in this region contain most individuals from each species and are comparatively younger than in the rest of the genome. The cross-coalescence times in the peak region are also significantly delayed, meaning that there is less recent gene flow than across other areas of the genome.

**Table 2:**
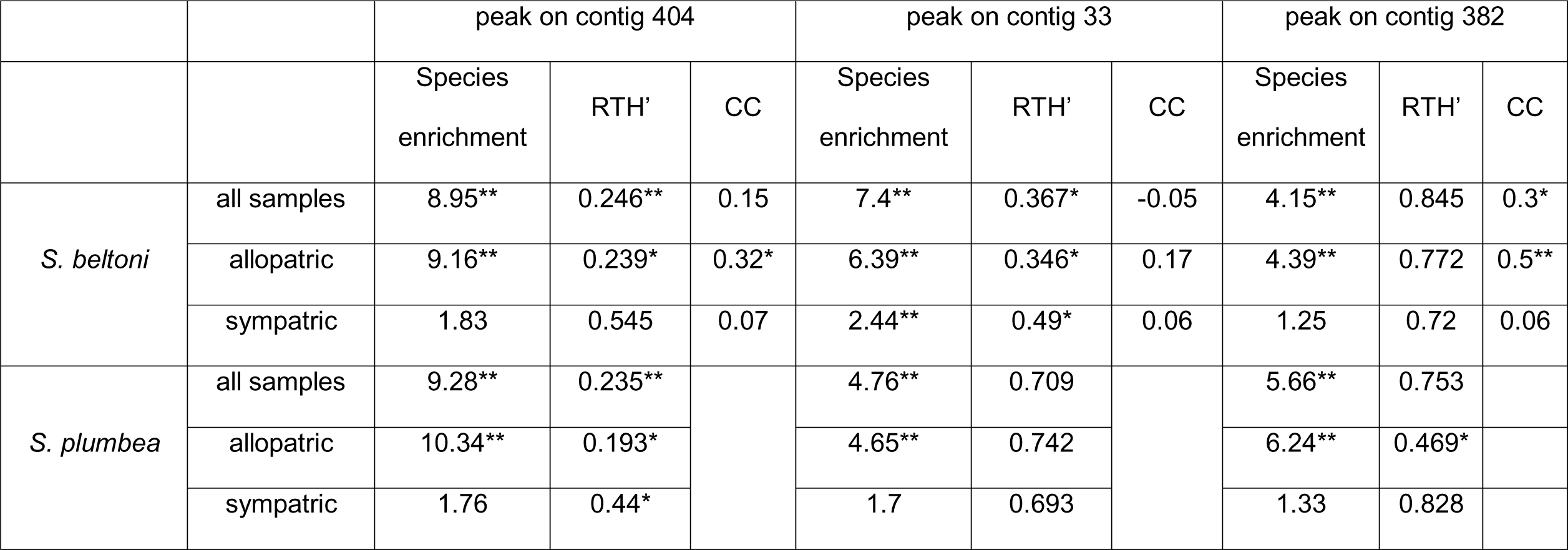
Results from statistical tests conducted on ARG-based statistics. Tests were conducted separately for allopatric, sympatric, or all samples combined. Species enrichment and RTH’ scores were compared to the distribution of these values across a set of control contigs without outlier peaks. Cross-coalescence (CC) values were compared between the peaks and their adjacent regions, and for control regions, between focal and adjacent areas. The distribution of the difference in CC values between focal and adjacent areas was assessed across all control regions and used to determine the statistical significance of the difference in CC values observed between the peak regions and those areas immediately adjacent to the peaks. Statistically significant results are shown by asterisks, which also denote the level of significance (*0.005; **0.001). Note that we ran species enrichment and RTH’ tests for each species, while the cross-coalescence test is run for the species pair and therefore only presented once in the top of the table.

## Discussion

Whole-genome sequencing of *Sporophila beltoni* and *Sporophila plumbea* revealed little genetic differentiation between this species pair despite their marked phenotypic differences, similar to what has been found in other *Sporophila* species [14,26,32]. The highest levels of differentiation were concentrated in three narrow peaks, containing only six genes, which are candidates for mediating the phenotypic differences between our focal taxa. However, unlike the other *Sporophila* species which have shown this pattern of genetic homogeneity [9,14], we inferred *S. beltoni* and *S. plumbea* to be comparatively old, in the order of 3 million generations (∼1 Myr). For comparison, the 10 species of southern capuchino seedeaters (those found south of the Amazon River [38]) began to diverge within a narrow window over the last million years [39]. Our demographic modelling suggests high levels of gene flow, and not recent origin, as a possible explanation for the genetic homogeneity between *S. beltoni* and *S. plumbea*. This scenario of relatively deep divergence, high overall gene flow, and a small number of highly differentiated regions suggests that these divergence peaks could be islands of speciation [40], i.e., regions of the genome that resist homogenization through gene flow. In contrast, in capuchino seedeaters most divergence peaks, which are also enriched for coloration genes, were found to be shaped through recent species-specific selective sweeps [9]. Therefore, divergence in regions of the genome containing pigmentation genes has likely evolved through different processes in *S. beltoni*/*plumbea* and capuchino seedeaters [9,14].

The two outlier peaks with the highest levels of differentiation and encompassing most of the highly differentiated SNPs (on contig 404 and contig 33), contained only three genes, two of which have functions that may mediate the phenotypic differences between our focal taxa. *EDN3* functions in melanoblast proliferation and differentiation, and mutations (including duplications) in either the *EDN3* gene itself or its receptors (*EDNR*s) can lead to hypo or hypermelanization in quails and chickens [41], and to several piebalding phenotypes in domestic pigeons [42]. These differences in pigmentation have been shown to affect not only feathers, but also other body parts, like the chicken comb [43] or even internal organs [41]. The Prolactin receptor (*PRLR*) is implicated in keratinization and feather formation [44] and Prolactin itself is known to stimulate molt [45] and is involved in coloration phenotypes in other taxa [46]. It is therefore likely that these genes are involved in mediating the coloration differences between *S. beltoni* and *S. plumbea*, the most notable of which is in the beak. These differences in beak color may contribute to promoting prezygotic reproductive isolation, as has been found between subspecies of Long-tailed Finches (*Poephila acuticauda*) [47,48]. Within the genus *Sporophila,* species have either gray, black, yellow or orange bills [24]. The clade containing *S. plumbea* and *S. beltoni* has a total of six species (also including *S. albogularis*, *S. falcirostris*, *S. collaris* and *S. schistacea*) [49], and four of these six species have orange bills, suggesting that the black melanized bill is possibly a derived trait in *S. plumbea*. More work is needed to understand if *EDN3* and *PRLR* mediate beak coloration across multiple *Sporophila* species, as is the case with the *BCO2* gene and carotenoid-based coloration in *Setophaga* warblers [50].

It was previously unclear whether *S. beltoni* and *S. plumbea* hybridized in their contact zone in Southern Brazil (Figure 1A, [34]), mainly due to the lack of individuals with a clear mixed phenotype in the wild. However, our study shows extensive admixture in this region, likely due to ongoing hybridization. Therefore, this narrow area of contact likely constitutes a hybrid zone. This hybrid zone coincides with a region where the breeding habitat of both species is degraded due to human activities, an aspect which may have contributed to increasing the overlap between the breeding ranges of both species and resulted in high levels of recent gene flow [51]. It remains to be determined whether this habitat alteration, and the possibly associated increase in gene flow, remains restricted to this narrow contact zone or if it may have consequences for the completion of speciation. Within the divergence peaks, sympatric individuals were more similar to each other than allopatric individuals, and in some analyses (e.g., PCA or admixture plots), some were indistinguishable (Figure 3B, Figure S1-S2). The exception was the peak on contig 33 which contained the *EDN3* gene that may be involved in controlling beak coloration, allowing the correct species identification in the contact zone (Figure 3B). Heterogeneous genomic landscapes tend to show a larger number of peaks outside of contact zones, where gene flow is reduced [52–54]. This increased level of divergence could be due to selection on traits in allopatry, unrelated to reproductive isolation and where gene flow does not occur [52]. If this is true for our focal taxa, the lack of admixture in *EDN3* in both allopatry and sympatry constitutes further evidence that this gene may mediate phenotypic differences relevant to reproductive isolation. However, we note that the genome-wide level of differentiation between *S. beltoni* and *S. plumbea* is very low in allopatry (F_ST_ = 0.0031), and that we infer high levels of genome-wide gene flow when analyzing sympatric or allopatric samples (Figure 2C). It is therefore possible that the three peaks of differentiation that we have identified have resisted gene-flow during speciation. This prompted us to search for signatures of different evolutionary processes which could have shaped our candidate genomic regions.

We did not find clear patterns using traditional summary statistics (e.g., Tajima’s D, nucleotide diversity, D_XY_) within the peak regions that would allow us to draw robust conclusions about the processes that have shaped them. We therefore leveraged the ARG and genealogy-based statistics that would potentially give us greater resolution and allow us to distinguish between two different extreme scenarios which could elevate F_ST_: recent and species-specific selective sweeps and resistance to gene flow [9,17]. Regions which have undergone recent species-specific selective sweeps are expected to show clades containing most individuals from the relevant species that are significantly shallower (i.e., showing reduced diversity) than other areas of the genome. Islands of speciation, on the other hand, should show signatures of reduced gene flow (i.e., delayed cross-coalescence) when compared to other regions of the genome. We note that these two extreme models are not mutually exclusive, and a locus could undergo a selective-sweep after which gene flow between species would be reduced [9]. This latter scenario is supported by our data in the peak regions, where we tend to see comparatively shallow clades for one or both species, as expected for loci that have undergone selective sweeps. Moreover, we also see evidence for delayed cross-coalescence (primarily outside of the hybrid zone, but also more generally when the p-value thresholds are not as stringent; Table 1 vs. Table S2). Taken together, our findings support the islands of speciation model [11,40], where a few loci that may be relevant to generating coloration differences and prezygotic isolation, have undergone selection and resisted gene flow.

Our study represents the first investigation into the phenotypic differences between *S. beltoni* and *S. plumbea* and shows how coloration genes can diverge early in speciation and likely mediate prezygotic isolation. We also exemplify how the ARG can be leveraged to distinguish between competing processes which shape genome evolution, finding evidence of reduced gene flow in speciation islands compared to the rest of the genome. Taken together, our genome scan, demographic reconstructions, and ARG-based analyses reveal the complicated interplay between selection and gene flow in shaping species boundaries.

## Materials and methods

### Sampling and dataset

We sampled a total of 36 individuals for whole genome resequencing, 18 *S. beltoni* and 18 *S. plumbea*. Samples originated from allopatric portions of the ranges of both species (n=11 for each species), as well as from the contact zone (n=7 for each species) (Figure 1; Table S1). For this study we only used adult males so that identification would be unambiguous, and so that our depth of coverage for autosomes and sex chromosomes would be equivalent. Birds were captured during breeding seasons (austral spring/summer) between 2008 and 2014 using mist nest, and subsequently banded and bled before being released. To conduct fieldwork in Brazil and collect samples, we secured licenses from the National Center for Research on the Conservation of Wild Birds (CEMAVE, licenses 3711 and 3778) and the Instituto Chico Mendes of Conservation of Brazilian Biodiversity (ICMBio), through the Brazilian System and Information on Biodiversity (SISBIO, license numbers: 13310, 36881, 35434, and 52714).

### Sequencing and variant discovery

We used the DNEasy blood and tissue kit (Qiagen, CA, USA) to obtain DNA from blood and prepared libraries following the protocol recommended for the TruSeq Nano DNA library preparation kit protocol (550 bp inset size). Libraries were pooled into two groups of 18 samples, using concentrations of adapter-ligated DNA (determined through digital polymerase chain reaction), and each group was sequenced on its own Illumina NextSeq 500 lane at the Cornell Institute for Biotechnology core facility. This produced a total of ∼1.6 billion 151 bp paired-end reads, and consequently based on the number of raw reads obtained for each individual, we expected an average depth of coverage of 5.6 +/- 1.1. We assessed sequence quality and performed filtering and adapter removal as described previously [17]. We first trimmed low-quality bases from individual reads and collapsed overlapping paired reads using AdapterRemoval version 2.1.1 (options: --trimns --trimqualities --minquality 10) [55]. We then aligned the filtered sequences from each individual to the *Sporophila hypoxantha* reference genome [14] using Bowtie2 version 2.4.3 [56] with the very sensitive local option. Despite using a reference genome from a different, yet closely related species, the average alignment rate was 96% (Table S1). The average depth of coverage across all samples and sites, calculated using Qualimap v2.2.1 [57], was 5.4 +/- 1.1x (Table S1). We followed the genotyping pipeline described in detail in [17] and filtered the resulting SNP dataset with VCFtools version 0.1.16 [58]. Briefly, we first marked PCR duplicates and realigned around indels using “MarkDuplicates” and “IndelRealigner”, respectively. We subsequently used the “Haplotypecaller” and “GenotypeGVCFs” modules from GATK version 3.8.1 [59] to identify variants and conduct joint genotyping across samples, resulting in a single variant file for the entire dataset. We then selected SNPs with the “SelectVariants” module of GATK (option: --selectType SNP) and filtered out those that did not satisfy the following filters: QD < 2, FS > 60.0, MQ < 30.0, ReadPosRankSum < -8.0. Finally, we performed additional post-hoc filtering using VCFtools to retain 15,663,387 variants present in at least 85% of individuals, with mean depth of coverage between 2 and 50 and a minor allele count of at least six. We explored the sensitivity of our dataset to a range of filtering parameters in different combinations, including up to a minimum depth of coverage of 5, a minor allele count of at least 10 and less that 5% missing data. These different filters produced congruent patterns in our PCAs and Manhattan plots, yet reduced the number of total SNPs retained. We therefore decided to retain a larger number of SNPs and, unless otherwise stated, use the dataset described above for downstream analyses.

### Analysis of mitochondrial DNA

Because we did not recover high-quality mitochondrial genomes for every individual from the whole genome data, possibly due to miss-assemblies related to nuclear sequences of mitochondrial origin, we decided to sequence the cytochrome c oxidase I (*COI*) gene, which is commonly used for species identification [60]. We used primers BirdF1 and COIBirdR2 and followed the procedures described by Kerr et al. [61]. We Sanger sequenced PCR-amplified DNA using both the forward and reverse primers at the Cornell Institute for Biotechnology core facility. Forward and reverse sequences were combined into a consensus sequence for each individual and aligned using Geneious version 10.2.6 [62]. In total we obtained 27 *COI* sequences, 8 allopatric and 6 sympatric *S. beltoni* and 7 allopatric and 6 sympatric *S. plumbea* (Table S1). We used these sequences to construct a *COI* haplotype network using the R version 4.0.2 [63] package pegas v1.1 [64].

### Genetic differentiation, admixture and summary statistics

We first assessed genome-wide patterns of differentiation among species and sampling locations (i.e., allopatric or sympatric) by conducting a Principal Component Analysis (PCA) in the R package SNPRelate version 3.3 [65]. Two samples originated from individuals that were later found to be related (TNN26 and TNN27, relatedness of 0.538) and separated from the remaining samples in a PCA (Figure S1A). We therefore excluded these samples from the PCA in Figure 1C, which produced the same pattern as when the samples were retained, and PC2 vs. PC3 were plotted (Figure S1B). We used the same programs to produce PCAs from the SNPs found within the divergence peaks, after subsetting the dataset by genomic location in VCFtools. We also calculated F_ST_ values for individual SNPs using VCFtools and displayed these in a histogram produced in R or as a Manhattan plot using the R package qqman version 0.1.8 [66]. For the latter, we simplified the plot by only visualizing SNPs with F_ST_ > 0.2. We also used VCFtools to subset the dataset by species and contig to calculate statistics base on the frequency of the most common haplotypes (H1, H2, H12 and H2/H1) in SelectionHapStats [67]. We used a window size (non-overlapping) of 25 SNPs and merged haplotypes differing in one position (-- distanceThreshold 1). We also used VCFtools to calculate R2 values and estimate linkage disequilibrium (LD) among all the different sites showing the highest differentiation (F_ST_>0.7) in the divergence peaks. For these same variants (24 on contig 404, 9 on contig 33 and 6 on contig 382), within each divergence peak, we explored local haplotypes by first phasing and imputing missing data using default parameters in BEAGLE version 3.3.2 [68]. We plotted the two haplotypes for each peak region from every individual using the function phylo.heatmap from the R package phytools [69], and clustered them according to similarity by producing a distance matrix in the R package vegan [70]. Finally, we explored ancestry along the contigs of interest by running Admixture version 1.3.0 [71] in 50 kb (non-overlapping) sliding windows using a K value of two. As the peak on contig 382 was narrow (∼2 kb), we used a sliding window of 10 kb in this case, as a compromise between not averaging out across too many SNPs while maintaining sufficient resolution. We plotted these values separately for allopatric and sympatric samples using a smoothing line in ggplot2 [72]. To calculate Tajima’s D and nucleotide diversity (π) in the contigs of interest, we used a vcf file that followed the same filtering steps described above except for that it was not filtered by minor allele frequency (the same file used in the demographic reconstruction), as this would bias both summary statistics that are sensitive to low frequency alleles. We calculated these statistics in non-overlapping, 5 kb windows. Finally, we calculated D_XY_ using the program pixy version 1.2.7 [73]. For this specific analysis we included invariant sites by running GATK’s “GenotypeGVCFs” module with the option “–includeNonVariantSite”. Following filtering recommendations [73], we then separated our data into variant and invariant sites and filtered them independently, as applying population genetic based filters such as minor-allele frequency (MAF) filtering would result in the loss of our invariant sites. We omitted the MAF filter to obtain invariant sites, but all other filtering steps were implemented as described previously (e.g., missing data or coverage filters).

### Demographic reconstruction

We estimated current and ancestral effective population sizes for both species, the splitting time between the taxa and migration rates (total of six demographic parameters) using an isolation with migration model as implemented in G-PhoCS version 1.3 [74]. Because this analysis is computationally intensive, we subsampled individuals from the dataset. We conducted three different analyses using individuals from different parts of the ranges of both species: allopatric (eight from each species), sympatric (seven from each species) or a combination of both types of samples (4 allopatric and 4 sympatric from each species). To avoid biasing our analyses by excluding low frequency alleles, we used a vcf file that was not filtered for minor allele frequency (∼61M SNPs), but otherwise had the same filters applied as described previously. We subsequently discarded contigs that were smaller than 1 Mb, Z-linked, or had outlier SNPs (contig 404, 33, 382 and 910). We used the “FastaAlternateReferenceMaker” module in GATK to generate sequence files for each individual for the remaining contigs, and sampled and aligned 2,500 loci, each 1 kb in length and at least 100 kb apart. We ran G-PhoCS for 1 million generations (for the sympatric and allopatric analyses) or for 300,000 iterations (for the combined analysis) and discarded the initial 50,000 as burn-in. We used the coda package in R [75] to check for convergence and subsequently converted median and 95% Bayesian credible intervals to generations or individuals (from mutation scale) by using an approximate mutation rate estimate of 10^-9^ per bp per generation [76]. We have focused our interpretations primarily on relative comparisons between the species which are independent of our assumption of mutation rate, as is the number of migrants per generation [74].

### Identification of genes in outlier genomic regions

We defined F_ST_ outlier regions as the genomic coordinates encompassed by the SNPs showing F_ST_ > 0.7. By using this approach, we aim to focus on the regions showing the highest levels of differentiation between species, however this does not imply that other regions, showing more subtle patterns, are not relevant to shaping phenotypic differences. We had previously mapped every contig in the reference genome to its corresponding chromosome in the Zebra Finch [14]. Most of these outlier SNPs were clustered together and located in three peaks (contig 404, contig 33 and contig 382), yet there was a single additional elevated SNP on scaffold 910 (near the gene LRRC15), corresponding to chromosome 9. We did not consider this a divergence peak as it included a single position, and don’t discuss it further. We searched for the genes within each divergence peak (+/- 5 kb) by inspecting our reference genome annotation using Geneious version 10.2.6 [62]. We used the NCBI database (http://www.ncbi.nlm.nih.gov/) to find information on the function of these different genes in other organisms. Finally, the outlier genomic regions were similar to the rest of the genome in average missing data (0.05 vs. 0.05), variant quality (QUAL; 1683.19 vs 1678.24) and depth of coverage (4.97 vs 5.00).

### Statistics derived from the Ancestral Recombination Graph (ARG)

We estimated local trees by inferring ARGs using the arg-sample module in ARGweaver version 1 [37]. We ran ARGweaver on unphased data in 5-7 mb intervals which included the areas of high differentiation in the three contigs with outlier peaks, plus a control set consisting of 10 contigs totaling ∼110 megabases. For the control set we broke large contigs into 5 mb fragments and ran the software following the settings described previously [17]. Briefly, we set the mutation and the recombination rates to 10−9/bp/gen, the effective population size to 500,000 individuals and the remaining parameters as follows: -c 5 -ntimes 20 -maxtime 1e7 -delta 0.005 -resample-window-iters 1 -resample-window 10000 -n 1000. We retained the last of 1,000 MCMC iterations, discarded the first and last 50 kb of each ARG block, and extracted trees every 500 bp. We used these trees to calculate three statistics as described in detail in [9] (Figure 4A). The species enrichment score measures the probability of observing subtrees of different sizes containing individuals from a certain species, and is calculated for both species and every subtree. For each tree we retained the maximum enrichment score obtained for a given species across all subtrees. A high enrichment score is obtained when a local tree has a clade with most or all individuals from that species. We note that when only two species are present in a tree like in this study, when one species enrichment score is high it means that individuals from the second species may also group together on the tree, and therefore both species enrichment scores will tend to be elevated. The relative TMRCA half-life [9,37], RTH’, is a normalized version of the time to most recent common ancestry (TMRCA). To calculate this statistic, we divided the time to the most recent common ancestor of half of the haploid samples for a species by the age of the youngest subtree containing at least half of all the haploid samples (irrespective of species). This normalization controls for the variation is coalescence times in different trees across the genome [37]. Finally, the inter-species cross coalescence measures the average age of the most recent coalescence events between tips in the tree which belong to the different species. We added the ages of the ten most recent cross-coalescence events in each tree and normalized them in the same way we did to calculate RTH’ [9]. We obtained these statistics for each tree (sampled every 500 bp) and then averaged those values across 5 kb non-overlapping windows. We generated empirical distributions for these statistics from the ∼22,000 windows (∼110 mb) obtained from the control contigs, and used these distributions to assess statistical significance as described previously [9,17]. The analysis was conducted separately for allopatric and sympatric samples, and a third time for all samples combined. We exported trees which show extreme enrichment scores for each species for illustration purposes. The code to calculate these three statistics and conduct these statistical tests is deposited in GitHub (https://github.com/CshlSiepelLab/bird_capuchino_analysis; see [9]).

## Acknowledgements

We thank Bronwyn Butcher for help with library preparation and sequencing. This project was funded through the Athena grant from the Cornell Lab of Ornithology (to T.N.N.) and NSF DEB-2232929 (to L.C.). The Brazilian Federal Agencies (CNPq and CAPES) provided scholarships to M.R. The Neotropical Grassland Conservancy (NGC) supported field work. The Brazilian National Research Council (CNPq grants (303318/2013, 309438/2016) supported C.S.F. We would like to acknowledge the property owners who kindly authorized our fieldwork. We thank Renata Grieco for permission to reproduce the bird illustrations in Figure 1. The sequence data generated for this project is archived on GenBank (Bio Project PRJNA382416).

## Author contributions

Designed project: T.N.N., M.R. and L.C. Performed fieldwork: M.R. and C.S.F. Obtained genomic data: T.N.N. Analyzed genomic data: T.N.N. and L.C. Supervised research: L.C. Wrote manuscript: T.N.N. and L.C., with edits from all authors.

## Declaration of interests

The authors declare no competing interests.

## Notes

### Competing Interest Statement

The authors have declared no competing interest.

