## Supplementary Files for "Genomic islands of speciation harbor genes underlying coloration differences in a pair of Neotropical seedeaters"

**Table S1. Sample and sequencing information.** Individuals marked with an asterisk represent samples with both whole genome and COI sequence data.

| Sample ID | Museum ID[a] | Species/Population | Locality/State | Latitude | Longitude | Reads mapped (%) | Depth of coverage[b] |
|---|---|---|---|---|---|---|---|
| TNN 1* | MCP 4728 | *S. beltoni* South | Ituim, RS | -28.615973° | -51.377815° | 96.28% | 5.76 |
| TNN 2 | MCP 4729 | *S. beltoni* South | Ituim, RS | -28.607678° | -51.373792° | 90.34% | 5.93 |
| TNN 3 | MCP 4730 | *S. beltoni* South | Foz Rio dos Touros, RS | -28.422111° | -50.519623° | 96.16% | 5.56 |
| TNN 4* | MCP 4733 | *S. beltoni* South | Jaquirana, RS | -28.873508° | -50.434351° | 95.74% | 5.95 |
| TNN 6* | MCP 2782 | *S. beltoni* South | Jaquirana, RS | -28.878927° | -50.444529° | 88.90% | 6.11 |
| TNN 7* | MCP 4731 | *S. beltoni* South | Criúva, RS | -28.900848° | -50.815279° | 93.25% | 3.66 |
| TNN 8* | MCP 4745 | *S. beltoni* South | Criuva, RS | -28.900842° | -50.815289° | 95.49% | 5.27 |
| TNN 9 | MCP 4756 | *S. beltoni* South | Capela do Caravaggio, RS | -28.185344° | -50.888842° | 86.96% | 5.67 |
| TNN 10* | MCP 3064 | *S. beltoni* South | Capela do Caravaggio, RS | -28.137003° | -50.902941° | 88.18% | 6.72 |
| TNN 13 | MCP 4735 | *S. beltoni* South | Estancia do Meio, SC | -28.314070° | -50.264668° | 93.07% | 5.45 |
| TNN 17* | MCP 4728 | *S. beltoni* South | Coxilha Rica, SC | -28.312059° | -50.289694° | 96.79% | 3.97 |
| TNN 19* | MCP 3625 | *S. beltoni* North | Fazenda Cuiabá, PR | -24.406128° | -50.043561° | 96.37% | 5.78 |
| TNN 20* | MCP 4739 | *S. beltoni* North | Faz. Trajano, PR | -24.409857° | -50.135690° | 96.49% | 5.64 |
| TNN 21* | MCP 2759 | *S. beltoni* North | Agrolandia, Santo Amaro, PR | -24.454667° | -50.254417° | 98.56% | 5.67 |
| TNN 22* | MCP 2761 | *S. beltoni* North | Faz. Estiva, PR | -24.731914° | -50.362885° | 97.05% | 4.80 |
| TNN 23 | MCP 2762 | *S. beltoni* North | Faz. Estiva, PR | -24.727778° | -50.366444° | 96.63% | 4.84 |
| TNN 24* | MCP 2763 | *S. beltoni* North | Faz. Estiva, PR | -24.726079° | -50.367548° | 98.18% | 5.78 |
| TNN 25 | MCP 2479 | *S. beltoni* North | Faz. Iberá, PR | -24.673611° | -50.331833° | 98.40% | 7.20 |
| TNN 26 | MCP 4740 | *S. plumbea* North | Dourado/Itirapina, SP | -22.248273° | -47.894860° | 97.56% | 5.35 |
| TNN 27 | MCP 4744 | *S. plumbea* North | Dourado/Itirapina, SP | -22.258553° | -47.902603° | 96.96% | 5.16 |
| TNN 29* | MCP 4764 | *S. plumbea* South | UPD, SP | -24.267293° | -49.209717° | 92.74% | 9.01 |
| TNN 30* | MCP 2764 | *S. plumbea* South | UPD, SP | -24.265639° | -49.211806° | 98.25% | 5.16 |
| TNN 31* | MCP 2765 | *S. plumbea* South | UPD, SP | -24.279722° | -49.211139° | 98.18% | 5.36 |
| TNN 33* | MCP 4774 | *S. plumbea* South | UPD, SP | -24.289805° | -49.220556° | 98.43% | 6.11 |
| TNN 35* | MCP 3624 | *S. plumbea* South | Fazenda Cuiabá, PR | -24.411536° | -50.048134° | 98.14% | 5.45 |
| TNN 36 | MCP 4741 | *S. plumbea* South | Joaquim Murtinho, PR | -24.402202° | -49.867375° | 98.55% | 5.37 |
| TNN 38 | MCP 4762 | *S. plumbea* South | Casa de Pedra, Faz 4N, PR | -24.388767° | -50.017134° | 98.73% | 6.16 |
| TNN 42 | MCP 4760 | *S. plumbea* North | Serra da Canastra, MG | -20.143074° | -46.860252° | 97.21% | 5.53 |
| TNN 45 | MCP 4737 | *S. plumbea* North | Parna Emas, GO | -18.262267° | -52.888989° | 96.84% | 5.36 |
| TNN 47 | MCP 4750 | *S. plumbea* North | Parna Emas, GO | -18.275788° | -52.855073° | 96.90% | 4.43 |

| | | | | | | | |
|---|---|---|---|---|---|---|---|
| TNN 50* | MCP 4775 | *S. plumbea* North | Campos do Encanto, MT | -14.997864° | -59.929392° | 96.90% | 4.26 |
| TNN 53* | MCP 4770 | *S. plumbea* North | PARNA C. Dos Guimarães, MT | -15.306970° | -55.815265° | 97.02% | 4.56 |
| TNN 58 | MCP 4743 | *S. plumbea* North | Pousada Cristal, MT | -15.798939° | -51.924395° | 97.76% | 2.48 |
| TNN 61* | MCP 4747 | *S. plumbea* North | Jalapão, TO | -10.568517° | -47.225987° | 96.92% | 4.52 |
| TNN 62* | MCP 4752 | *S. plumbea* North | Jalapão, TO | -10.289050° | -46.941939° | 97.51% | 4.96 |
| TNN 65* | MCP 4757 | *S. plumbea* North | PN Grande Sertão Veredas, MG | -15.252919° | -45.634753° | 97.38% | 6.13 |
| TNN 16 | MCP 4734 | *S. beltoni* South | Coxilha Rica, Lages, SC | -28.312019° | -50.289409° | COI only | |
| TNN 18 | MCP 4759 | *S. beltoni* North | Fazenda Cuiabá, Piraí do Sul, PR | -24.403965° | -50.041055° | COI only | |
| TNN 37 | MCP 4772 | *S. plumbea* South | Fazenda 4N, Piraí do Sul, PR | -24.384040° | -50.022276° | COI only | |
| TNN 49 | MCP 4766 | *S. plumbea* North | Parna Emas, GO | -18.305292° | -52.903956° | COI only | |
| TNN 55 | MCP 4753 | *S. plumbea* North | Chapada dos Guimarães, MT | -15.342566° | -55.786049° | COI only | |

[a]MCP: Pontificia Universidade Catolica do Rio Grande do Sul, Museu de Ciencias e Tecnologia.
[b]Calculated from BAM files using Qualimap v2.2.1.

**Table S2: Statistical tests for ARG-based statistics**. This table differs from the equivalent table in the main text in that it also shows the two less stringent significance cut offs (*0.05; **0.01; ***0.005; ****0.001). Other details as in Table 2.

| | | peak on contig 404 | | | peak on contig 33 | | | peak on contig 382 | | |
|---|---|---|---|---|---|---|---|---|---|---|
| | | Species enrichment | RTH’ | CC | Species enrichment | RTH’ | CC | Species enrichment | RTH’ | CC |
| *S. beltoni* | all samples | 8.95**** | 0.246**** | 0.15 | 7.4**** | 0.367*** | -0.05 | 4.15**** | 0.845 | 0.3*** |
| | allopatric | 9.16**** | 0.239*** | 0.32*** | 6.39**** | 0.346*** | 0.17* | 4.39**** | 0.772* | 0.5**** |
| | sympatric | 1.83* | 0.545** | 0.07 | 2.44**** | 0.49*** | 0.06 | 1.25 | 0.72* | 0.06 |
| *S. plumbea* | all samples | 9.28**** | 0.235**** | | 4.76**** | 0.709** | | 5.66**** | 0.753* | |
| | allopatric | 10.34**** | 0.193*** | | 4.65**** | 0.742* | | 6.24**** | 0.469*** | |
| | sympatric | 1.76* | 0.44*** | | 1.7* | 0.693* | | 1.33 | 0.828 | |

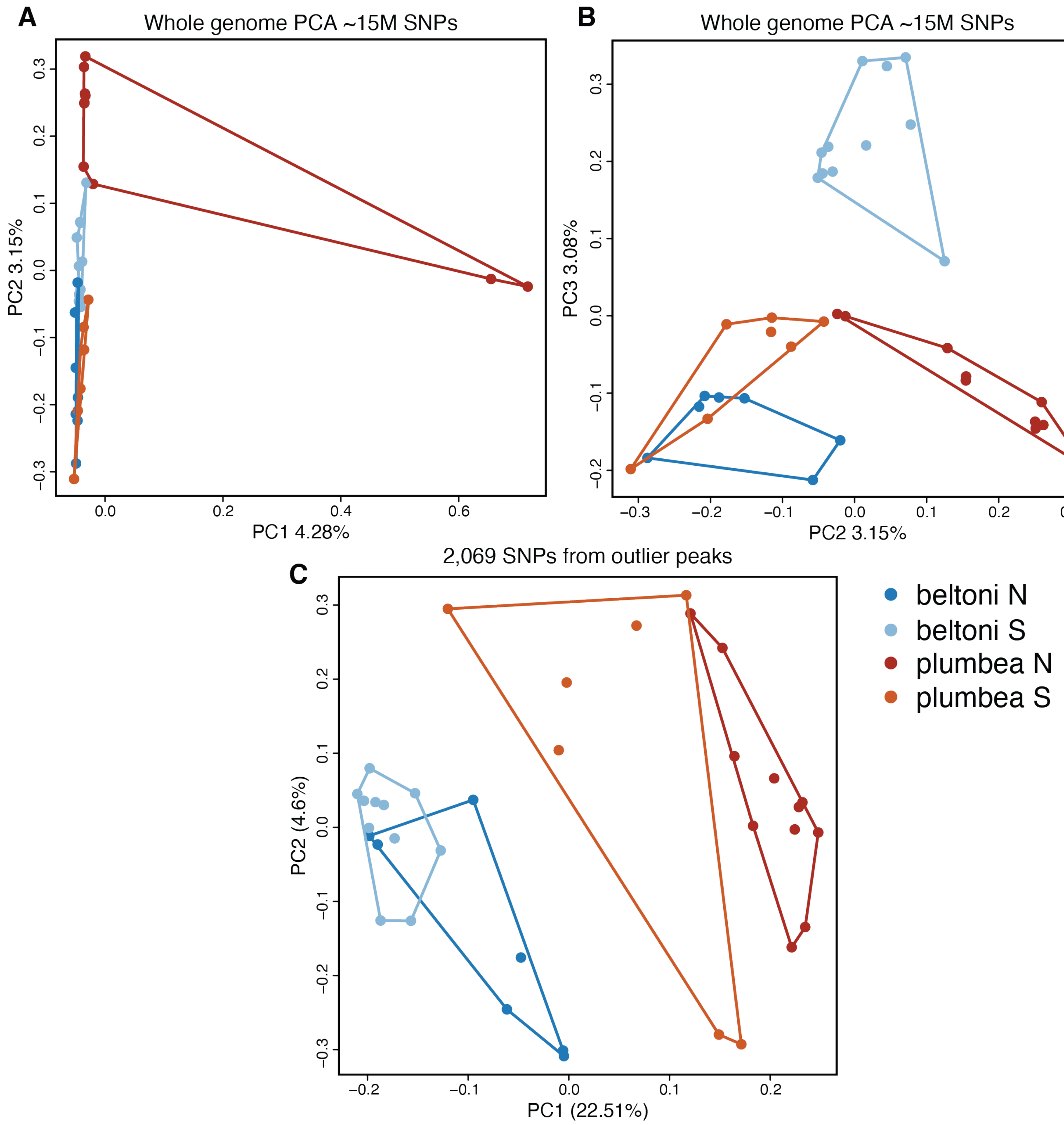


**Figure S1: A.** Principal Component Analysis (PCA) derived from ~15 million genome-wide SNPs (the same data as in Figure 1C) without excluding samples TNN26 and TNN27 from Itirapina, SP. These samples are from related individuals (relatedness of 0.538) and separate from the rest of the dataset in PC1. **B.** The same PCA as in **A** but this plot shows PC2 vs. PC3 and displays the same pattern as Figure 1C (where TNN26 and TNN27 were excluded). **C.** PCA derived from 2,069 SNPs from within the outlier regions of contigs 404, 33, and 382. Samples clustered by species within these peaks of divergence, despite low overall genomic differentiation.

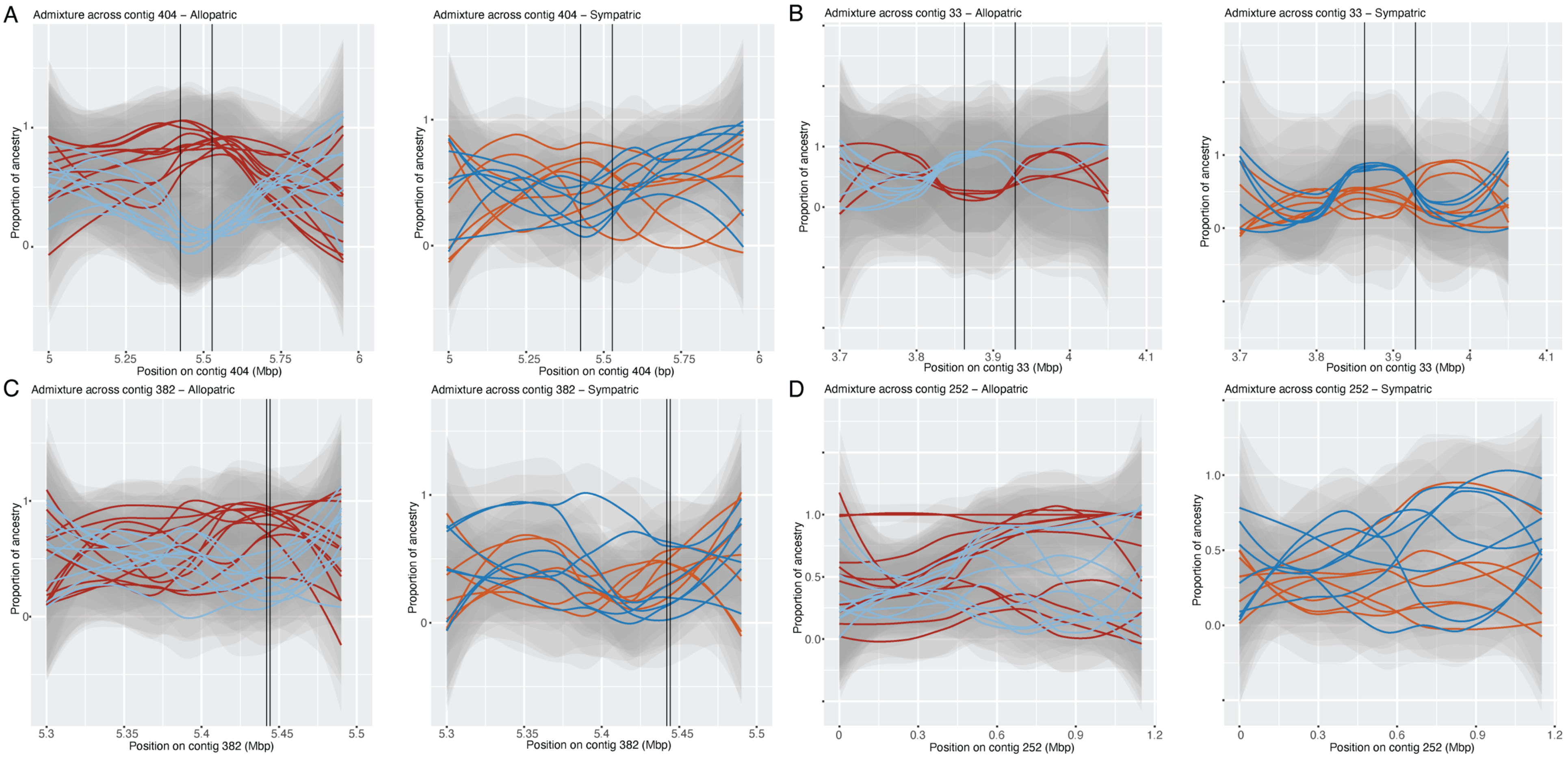


**Figure S2: Sliding-window ancestry values**. We calculated ancestry in Admixture in 50 kb sliding windows (except for contig 382 where we used 10 kb windows because the peak was narrow) and plotted values for each individual using a smoothing line. Outlier regions on contig 404 (**A**), 33 (**B**), and 382 (**C**) are indicated between vertical lines. Contig 252 (**D**) is a control contig without an outlier region. We plotted allopatric (on the left) and sympatric (on the right) samples separately. For allopatric samples, the plots show increased separation between individuals in the peak regions, and mixed ancestry immediately outside of these areas. For sympatric samples, we observed separation between individuals only in the peak on contig 33 (**B**). The control contig (**D**) shows admixed individuals and little resolution regardless of the contig coordinate and sample origin. Ancestry ranges from 0 to 1, but the smoothing algorithm makes the plots extend beyond this range because of the uncertainty shown by the confidence bands, yet we note that values beyond the [0,1] interval should not be interpreted.

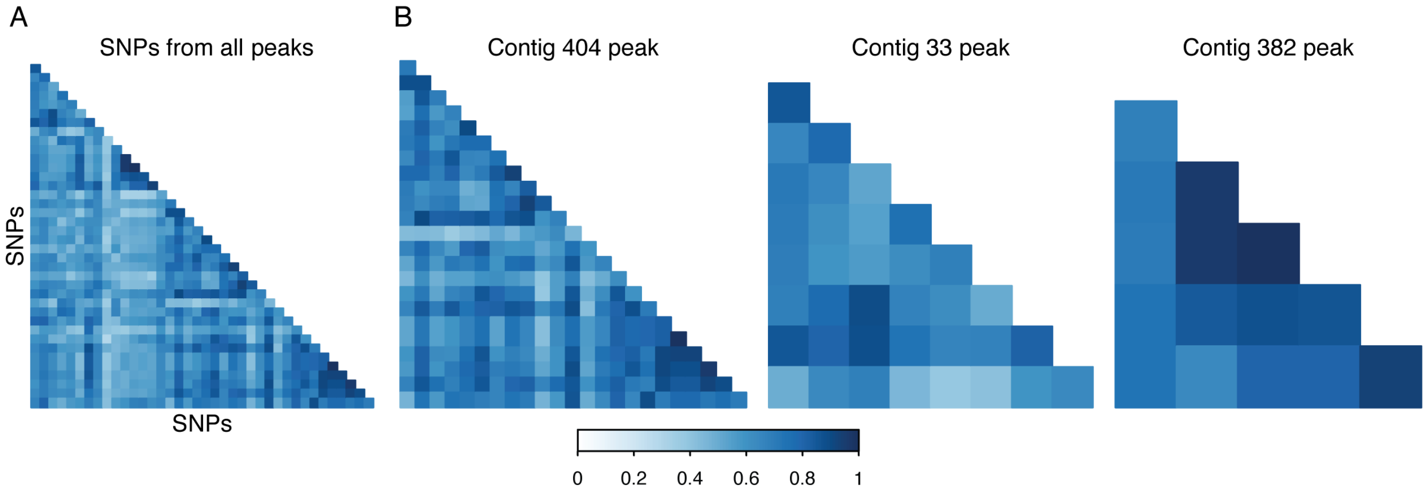


**Figure S3: Linkage disequilibrium among the SNPs showing the highest $F_{ST}$ values.** LD measured by the r2 statistic among all combinations of the 39 SNPs with $F_{ST} > 0.7$ from the peak regions on contig 404, 33 and 382 (note that these contigs belong to different chromosomes). The three plots on the right show within-peak comparisons (24 SNPs for the peak on contig 404, 9 SNPs for contig 33, and 6 SNPs for contig 382). The average r2 statistic obtained when comparing SNPs within peaks was only slightly higher than when comparing positions amongst peaks and therefore chromosomes (average intra-chromosomal r2: 0.7, range: 0.36-1; average inter-chromosomal r2: 0.57, range: 0.29-0.89). The average r2 values within contig were similar for contig 404 (0.7, range: 0.36-1), contig 33 (0.66, range: 0.38-0.9), and contig 382 (0.82, range: 0.65-1).

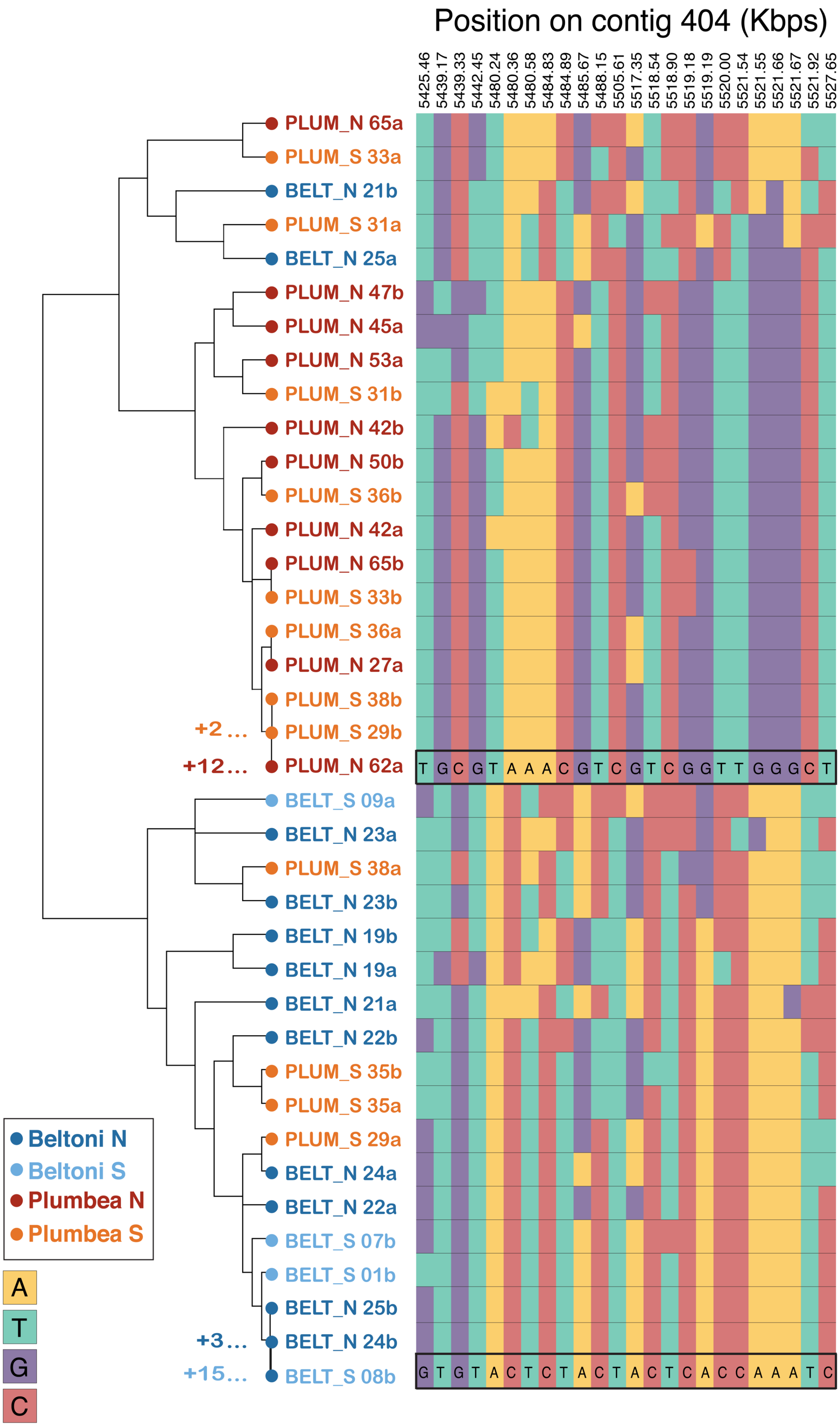
Position on contig 404 (Kbps)
5425.46
5439.17
5439.33
5442.45
5480.24
5480.36
5480.58
5484.83
5484.89
5485.67
5488.15
5505.61
5517.35
5518.54
5518.90
5519.18
5519.19
5520.00
5521.54
5521.55
5521.66
5521.67
5521.92
5527.65
PLUM_N 65a
PLUM_S 33a
BELT_N 21b
PLUM_S 31a
BELT_N 25a
PLUM_N 47b
PLUM_N 45a
PLUM_N 53a
PLUM_S 31b
PLUM_N 42b
PLUM_N 50b
PLUM_S 36b
PLUM_N 42a
PLUM_N 65b
PLUM_S 33b
PLUM_S 36a
PLUM_N 27a
PLUM_S 38b
+2 ...
PLUM_S 29b
+12 ...
PLUM_N 62a
T G C G T A A A C G T C G T C G G T T G G G C T
BELT_S 09a
BELT_N 23a
PLUM_S 38a
BELT_N 23b
BELT_N 19b
BELT_N 19a
BELT_N 21a
BELT_N 22b
PLUM_S 35b
PLUM_S 35a
PLUM_S 29a
BELT_N 24a
BELT_N 22a
BELT_S 07b
BELT_S 01b
BELT_N 25b
+3 ...
BELT_N 24b
+15 ...
BELT_S 08b
G T G T A C T C T A C T A C T C A C C A A A T C
Beltoni N
Beltoni S
Plumbea N
Plumbea S
A
T
G
C

**Figure S4: Haplotypes from the peak region of contig 404**. Phased genotypes for the 24 SNPs showing high differentiation ($F_{ST}$ > 0.70) in the peak region of contig 404. Each individual is represented by its two haplotypes (labeled a and b), which are color-coded by species and geographic origin. Individuals are labeled with their species (BELT for *S. beltoni* or PLUM for *S. plumbea*), geographic origin (North or South) and individual ID. Haplotypes are clustered in the tree on the left by their similarity, and the genotypes for the 24 sites are shown on the right, with the four nucleotides color-coded. Each species shows a common haplotype which is shown with a black box (note that for simplicity these were trimmed from the bottom of the two main clusters in the tree, and the number of haplotypes which were omitted are indicated on the left). There is comparatively less variation within species in each of these haplotypes than between species. Only a few individuals from the contact zone (*S. beltoni* North or *S. plumbea* South) do not possess two copies of the most common haplotype for their species. Of these, there are two heterozygote *S. beltoni* birds from the North (21 and 25) and two heterozygote *S. plumbea* birds from the South (29 and 38). Additionally, there is an *S. plumbea* individual from the South (35) which is a homozygote for the *S. beltoni* haplotype.

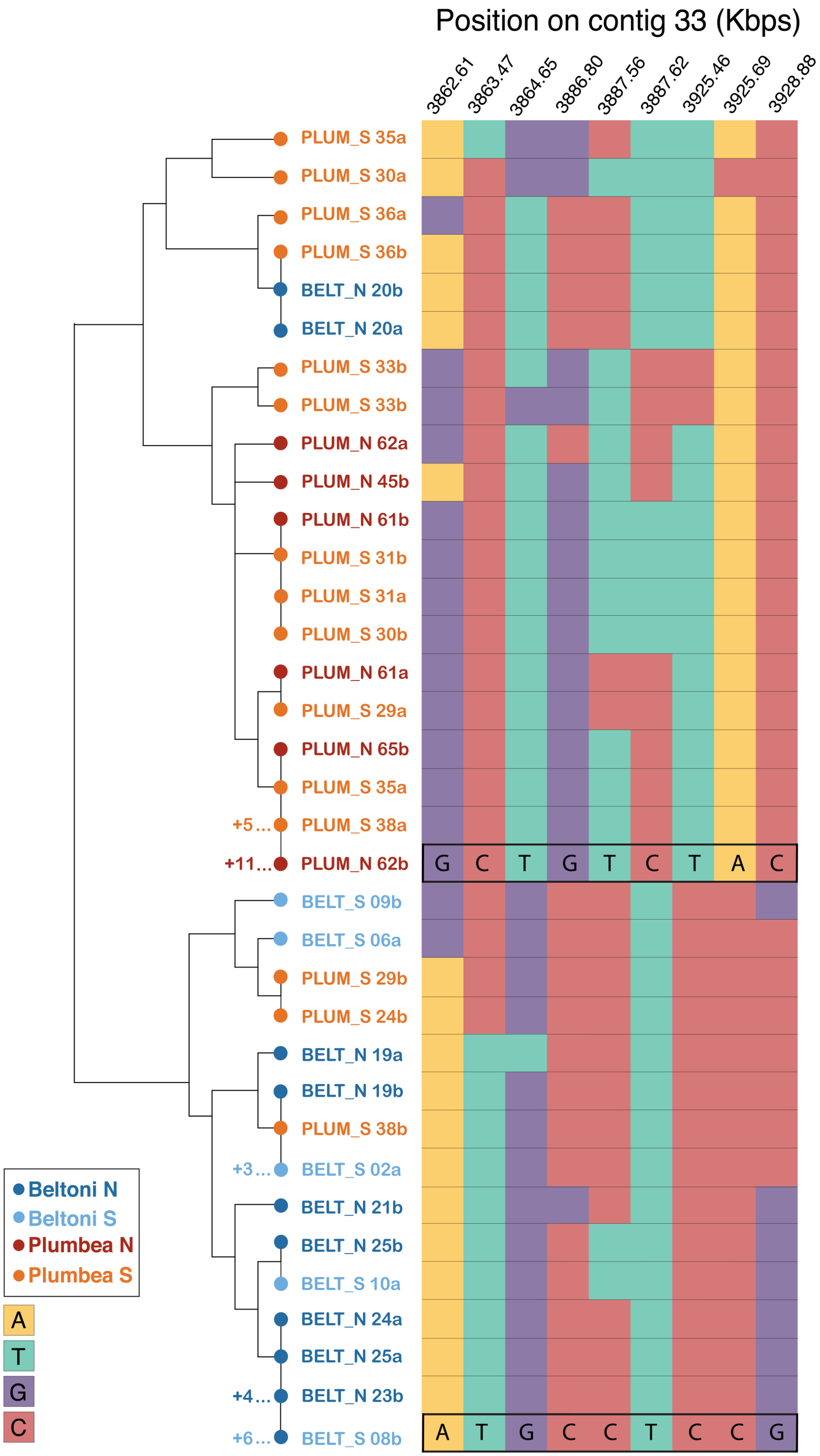


**Figure S5: Haplotypes from the peak region of contig 33.** Details as in Figure S4. Note that there is one Northern *S. beltoni* bird (20) that is a homozygote for the *S. plumbea* haplotype. Additionally, there are three Southern *S. plumbea* individuals that are heterozygotes (24, 29, 38).

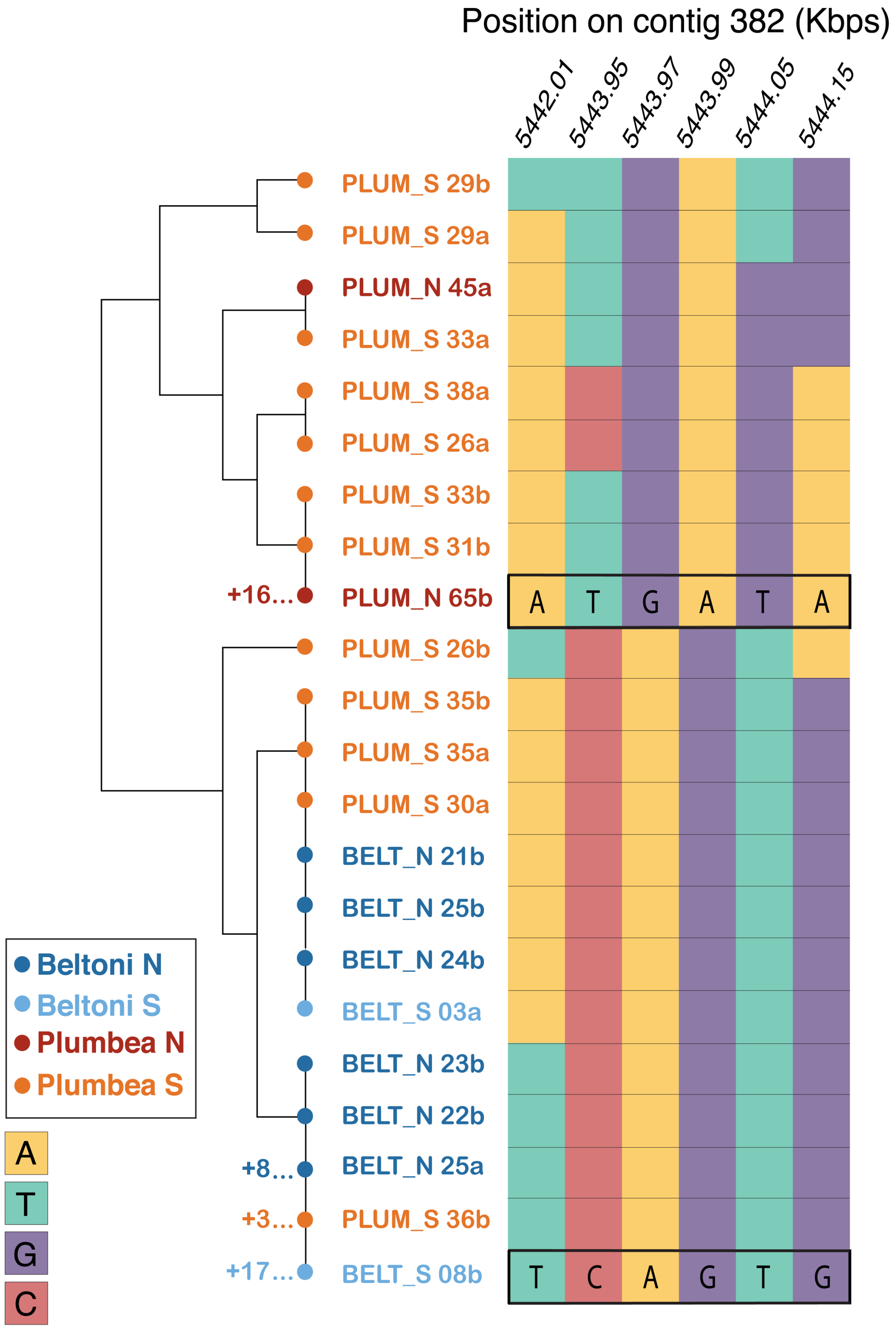


**Figure S6: Haplotypes from the peak region of contig 382.** Details as in Figure S4. Note that there are three Southern *S. plumbea* individuals that are heterozygous for the *S. beltoni* haplotype (26, 30, 36) and one individual that is homozygous (35).

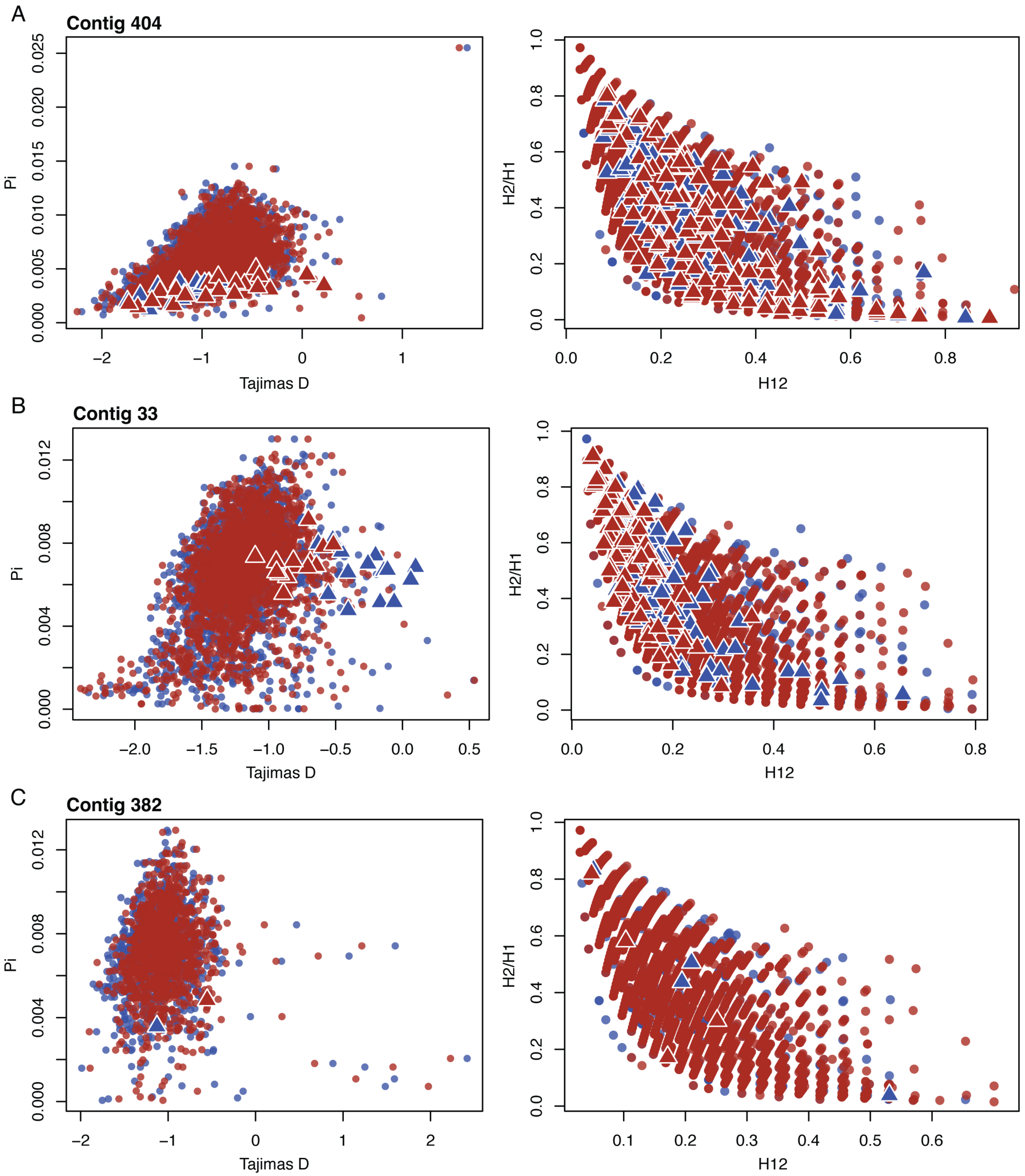


**Figure S7: Plots of summary statistics**. Biplots of Tajima's D vs. nucleotide diversity (left panels) and H12 vs. H2/H1 (right panels) for the contigs with peaks. Circles represent statistics derived from 5 kb windows (left) and 25-SNP windows (right), and are color coded by species (red: *S. plumbea*; blue: *S. beltoni*). The triangles represent windows that belong to the outlier peak region.

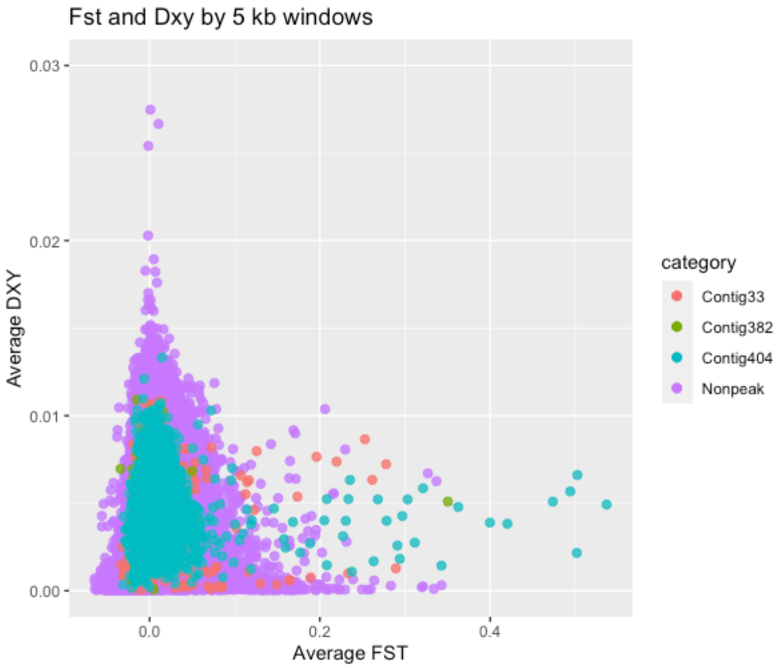


**Figure S8: Biplot of $F_{ST}$ and $D_{XY}$.** Average statistics for 5-kb windows. Contigs with and without outlier $F_{ST}$ peaks are color-coded. For simplicity we did not plot seven windows with outlier $D_{XY}$ values that were not in the $F_{ST}$ peak regions.

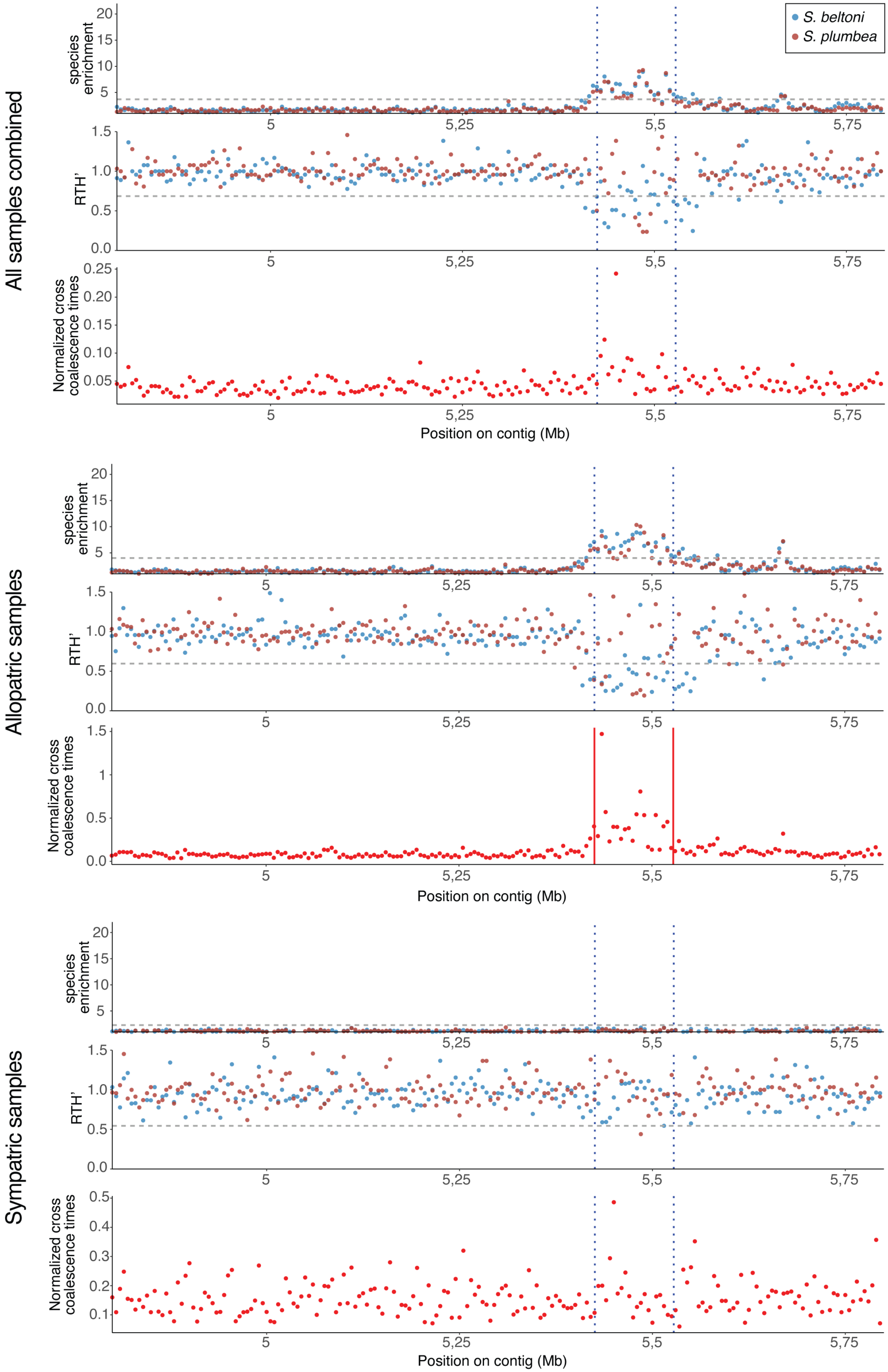

All samples combined
Allopatric samples
Sympatric samples
species enrichment
RTH'
Normalized cross coalescence times
S. beltoni
S. plumbea
Position on contig (Mb)
5
5,25
5,5
5,75

**Figure S9: ARG-derived statistics for contig 404**. Plots showing species enrichment values for 5 kb windows for *S. plumbea* and *S. beltoni* (top, color-coded), RTH' values for each species (middle), and cross-coalescence times for the species pair (bottom). Each set of three plots belongs to analyses done with all samples combined (top), allopatric samples only (middle), or sympatric samples only (bottom). The horizontal dashed lines in the species enrichment and RTH' plots represent the genome-wide threshold of statistical significance ($p<0.001$ for enrichment values and $p<0.005$ for RTH'). The thresholds are slightly different for each species, but for illustration purposes we just show one. For more detailed results see Table 2. Vertical dashed lines show the peak region. The solid red vertical lines show that the difference in cross-coalescence in the peak region compared to the adjacent area is statistically significant when compared to a distribution of this statistic obtained from control scaffolds.

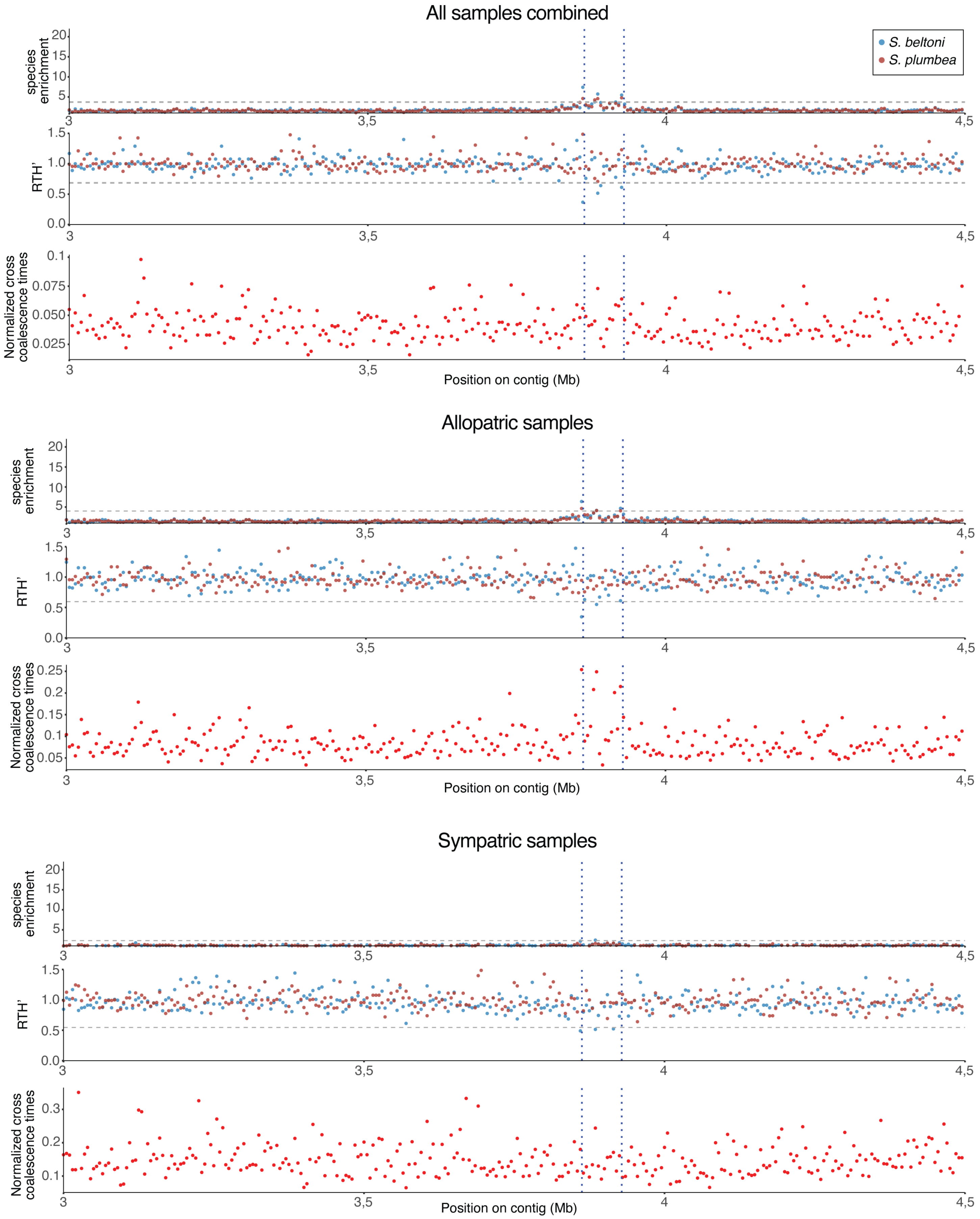


**Figure S10: ARG-derived statistics for contig 33.** Other details as in Figure S9.

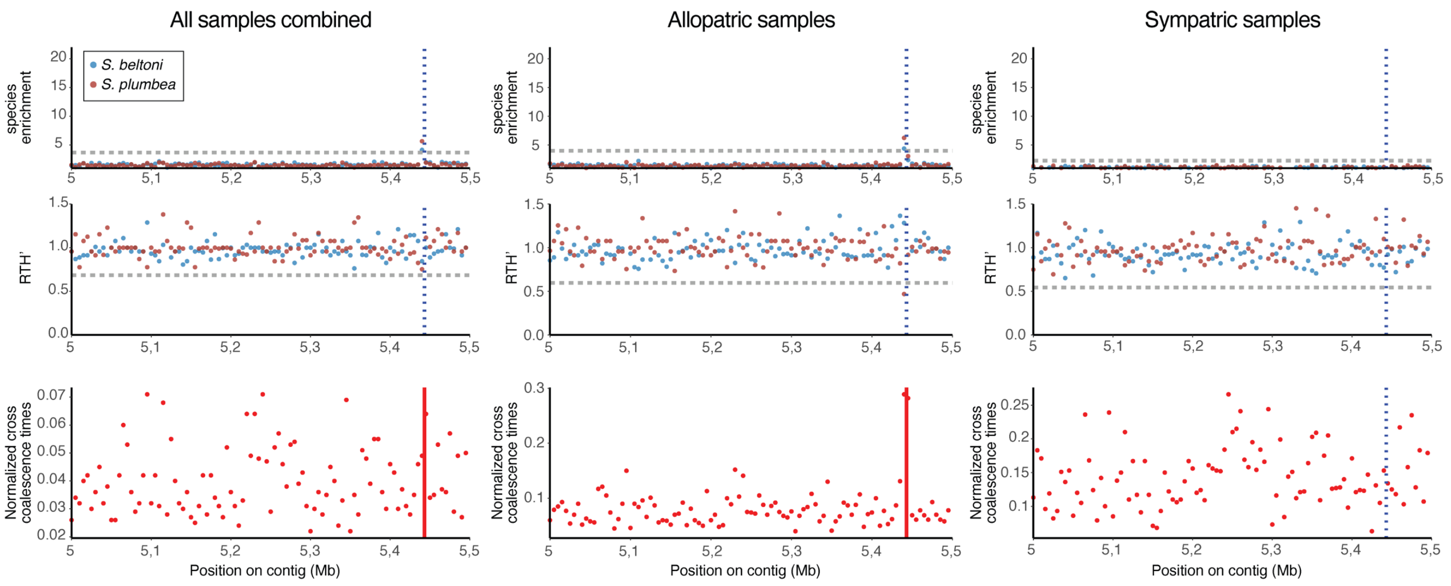


**Figure S11: ARG-derived statistics for contig 382.** Other details as in Figure S9.

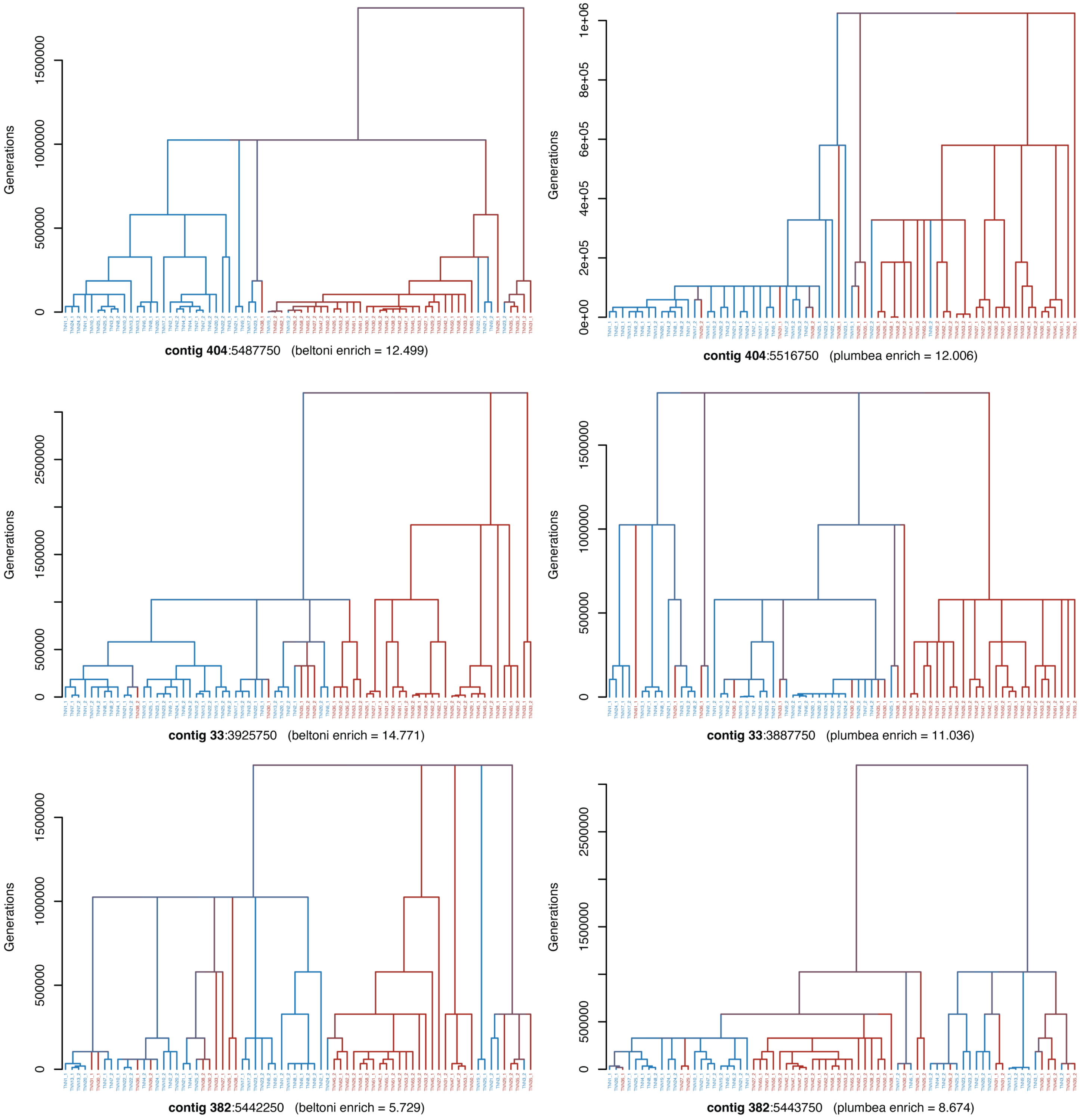


**Figure S12: Topologies obtained from the outlier peaks.** The trees represent the position within the peak of each contig (contig 404 on the top, contig 33 in the middle, and contig 382 in the bottom) which maximizes the species enrichment statistic (plots for *S. beltoni* are on the left and for *S. plumbea* are on the right). Each terminal represents the haploid genome from a sample, and these are color-coded in blue for *S. beltoni* and in red for *S. plumbea*. The color of internal branches is an average over all offspring branches. For each tree there is information of position on the contig in base-pairs, followed by the species enrichment which is maximized.
